# The Similarity Structure of Real-World Memories

**DOI:** 10.1101/2021.01.28.428278

**Authors:** Tyler M. Tomita, Morgan D. Barense, Christopher J. Honey

**Affiliations:** Department of Psychological & Brain Sciences, Johns Hopkins University, Baltimore, MD, USA; Department of Psychology, University of Toronto, Toronto ON, Canada; Rotman Research Institute, Baycrest Hospital, Toronto ON, Canada

## Abstract

How do we mentally organize our memories of life events? Two episodes may be connected because they share a similar location, time period, activity, spatial environment, or social and emotional content. However, we lack an understanding of how each of these dimensions contributes to the perceived similarity of two life memories. We addressed this question with a data-driven approach, eliciting pairs of real-life memories from participants. Participants annotated the social, purposive, spatial, temporal, and emotional characteristics of their memories. We found that the overall similarity of memories was influenced by all of these factors, but to very different extents. Emotional features were the most consistent single predictor of overall memory similarity. Memories with different emotional tone were reliably perceived to be dissimilar, even when they occurred at similar times and places and involved similar people; conversely, memories with a shared emotional tone were perceived as similar even when they occurred at different times and places, and involved different people. A predictive model explained over half of the variance in memory similarity, using only information about (i) the emotional properties of events and (ii) the primary action or purpose of events. Emotional features may make an outsized contribution to event similarity because they provide compact summaries of an event’s goals and self-related outcomes, which are critical information for future planning and decision making. Thus, in order to understand and improve real-world memory function, we must account for the strong influence of emotional and purposive information on memory organization and memory search.

**Significance:** Our brains enable us to understand and act within the present, informed by previous, related life experience. But how are our life experiences organized so that one event can be related to another? Theories have suggested that we use spatiotemporal, social, causal, purposive, and emotional dimensions to inter-relate our memories; however, these organizing principles are usually studied using impersonal laboratory stimuli. Here, we mapped and modeled the connections between people’s own annotated life memories. We found that life events are linked by a variety of factors, but are predominantly connected in memory by their primary activity and emotional character. This highlights a need for theories of memory organization and retrieval to better account for the role of high-level actions and emotions.

## Introduction

Our memory function depends on the similarity structure of the information we encode and retrieve. Because we organize memories according to their similarities, a person who is asked which painting they saw at a gallery the previous week might begin by recalling a distinctive red painting, and then another painting with a red palette; or if they first recall a painting in a specific high-ceilinged room, they might next recall another painting from that same room. More generally, processes of priming, association, interference, and forgetting all depend on the similarity relationships between items in memory [Gonnerman et al., 2007, Ritvo et al., 2019]. Once we have mapped the similarity structure of information in memory [Shepard, 1980, Tversky, 1977], we can build quantitative models of how we search and sample from that space [Anderson and Bower, 1972, Gillund and Shiffrin, 1984, Howard and Kahana, 2002, Naim et al., 2020, Polyn et al., 2009]. However, we currently know little about the similarity structure of the memories that matter most to us — the events of our own lives, which shape our identities and our life decisions [Bluck, 2003, Bluck et al., 2005, Conway and Pleydell-Pearce, 2000].

What determines the similarity between two events in our lives? Separate strands of research have suggested that each of the basic dimensions we use to describe a scenario (“who,” “why,” “what,” “where,” and “when”) are important for relating memories. In social and person-based theories, agents and their traits are hubs for linking memories [Hastie, 1988, Srull and Wyer, 1989]. In script and schema theories, the main action sequence or activity, such as “dining at a restaurant”, is the central feature of a life memory [Kolodner, 1983, Reiser et al., 1985, Schank, 1983]. In the behavioral and systems neuroscience literatures, centered on hippocampal coding theories, the spatial environment and spatiotemporal coordinates provide an indexing system for memories [Maguire and Mullally, 2013, Nielson et al., 2015, O’Keefe and Nadel, 1978, Robin et al., 2018]. In human autobiographical memory literature, life memories are thought to be organized by their narrative structure, grouped according to causally-linked “event clusters” [Brown and Schopflocher, 1998, Conway and Pleydell-Pearce, 2000]. Finally, emotional valence and personal relevance have also been proposed as important and interacting variables that connect our life memories [Talmi, 2013, Wright and Nunn, 2000].

Because life memories are complex and high-dimensional, it is understandable that diverse organizational theories and principles have been proposed. However, because these strains of literature have proceeded somewhat independently, individual studies have tended to examine only one or two dimensions, as motivated by each independent theory. Therefore we lack a comparison of how these different memory dimensions combine to structure our life memory. Consider, for example, a young woman’s memory of playing soccer in the park with friends. Next, imagine changing individual dimensions of this memory – the location, the time, the primary activity, the participants, or the emotional character of the event (Figure 1). In each case, the altered memory remains related to the original memory, but the similarity of the two memories differs depending on which dimensions are varied. Which dimensions most strongly determine the similarity of our memories for complex real-world events? Here, we set out to fill this gap, characterizing the similarity structure of people’s real world memories, and the role of multiple dimensions in determining that similarity structure.

**Figure 1:**
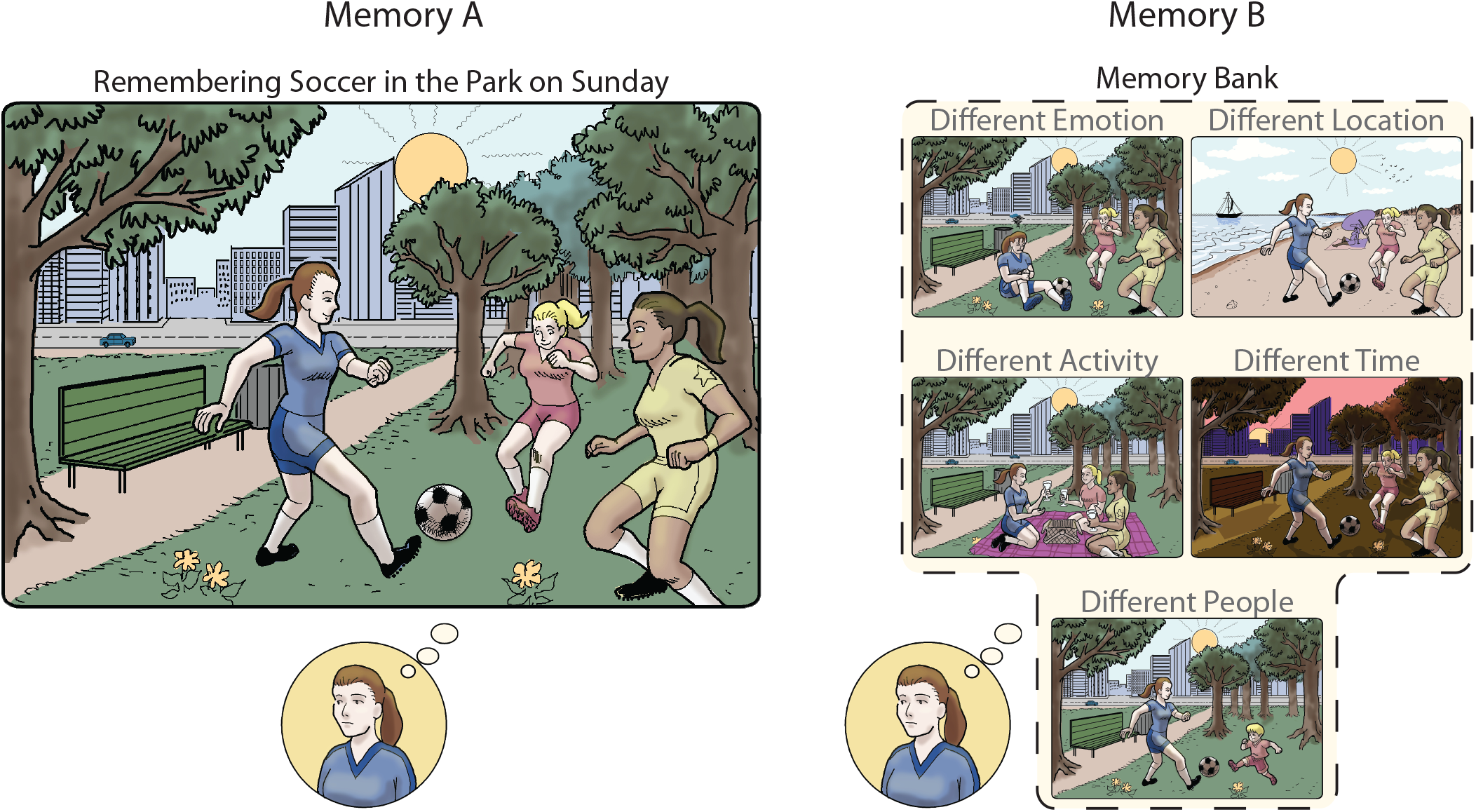
Which features determine the perceived similarity of real-world memories? This question is important because the similarity structure of our memories may guide memory search, and it may shape our identities by influencing how we think of one life event in the context of another. In this illustration, a woman is remembering playing soccer with two friends in the park the past Sunday. This is her original memory. Imagine that she has five other memories that are similar to this base memory, differing along only a single dimension: location, time, social setting, activity, or emotion. Which of these other memories are perceived as most (or least) similar to her original memory?

We mapped the links between memories using an “event-cueing” approach [Brown and Schopflocher, 1998]. Participants first recalled one life memory, and then a second life memory (Figure 2). Thereafter, participants annotated the memories and their inter-relationships. Our approach differs from prior studies of memory similarity in four ways. First, we employed a more theory-neutral and data-driven approach, measuring pairwise distance along 68 dimensions spanning spatiotemporal, social, emotional, and purposive features of memory. Second, our participant pool is larger and more diverse than prior literature on the structure of memory, which commonly employed undergraduates. Third, we analyzed the data using both descriptive statistics and predictive modeling, to demonstrate the generalizability of of our findings across participants. Fourth, in addition to the standard event cueing paradigm, we also investigated retrieval under conditions when participants were explicitly searching for memories that are very similar to or very different from a cue memory.

**Figure 2:**
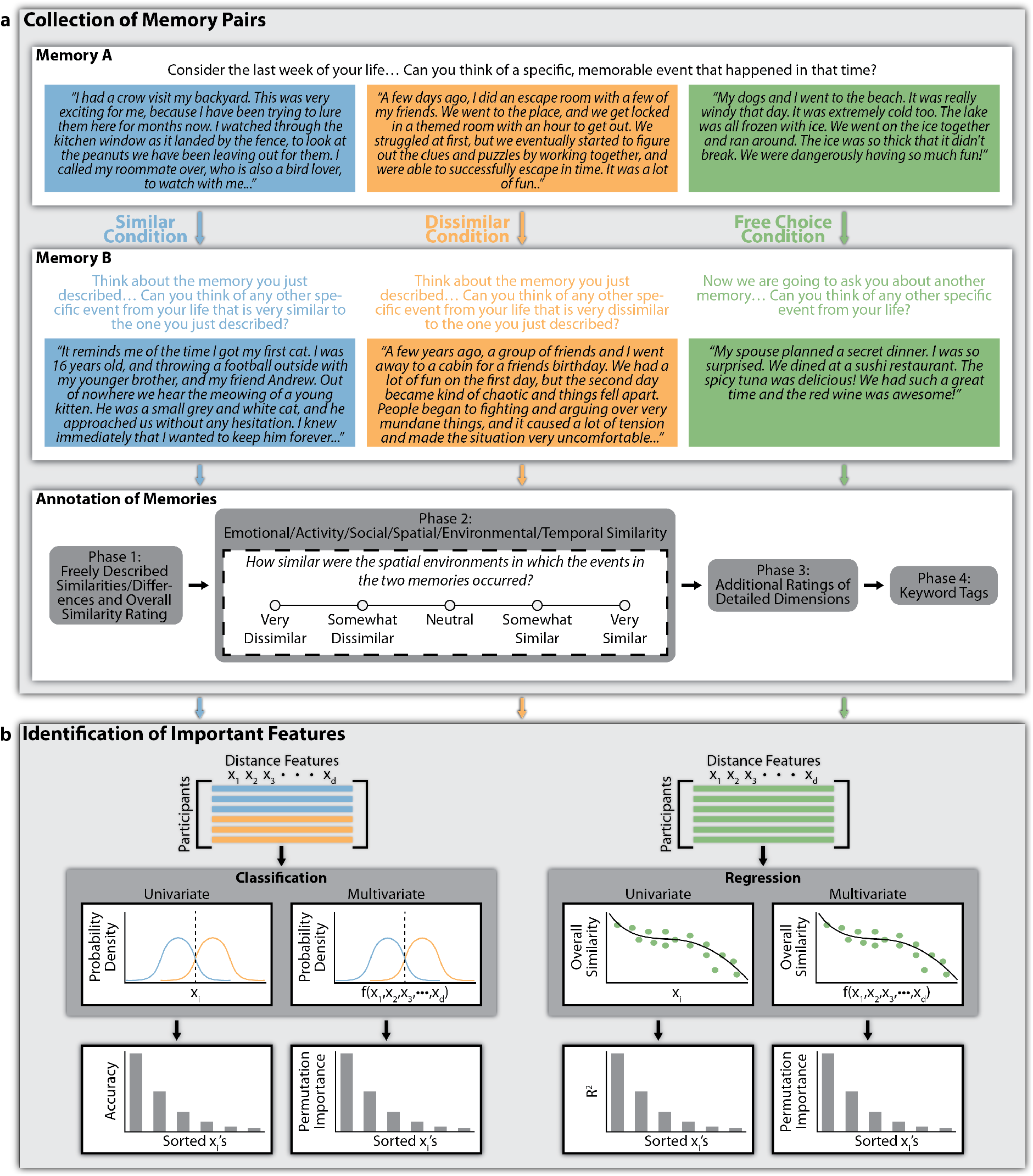
Collection of Annotated Memory Pairs and Identification of Features Driving Similarity Judgments. **(a):** Memory pairs were elicited from participants (“Memory A” and “Memory B” subpanels) who then annotated their memories (“Annotation of Memories” subpanel). Each participant was assigned to one of three conditions: Similar, Dissimilar, and Free Choice. In Phase 1 of annotation, participants were asked to describe the similarities and/or differences between the two memories, and rate their overall perceived similarity. In Phase 2, the similarity of the two memories along various dimensions was rated (e.g. emotional similarity, temporal proximity). In Phase 3, a variety of additional ratings were made on each of the two memories separately. In Phase 4, participants tagged each memory with one to five keywords of their choosing. **(b):** Distance features, each representing distance between the two memories along a different dimension, were derived from the annotations provided by each participant in Phase 2 and Phase 3. Using these distance features (each denoted by xi), along with explicit overall memory similarity judgments made for each memory pair, we examined which distance features had the strongest tendency to take on larger values for memory pairs judged to be dissimilar compared to memory pairs judged to be similar; such distance features should be the most predictive of similarity judgments. Thus we performed classification analysis (distinguishing Similar vs. Dissimilar conditions) and regression analysis (prediction of Overall Similarity ratings in the Free Choice condition) to identify individual features (univariate) and combinations of features (multivariate) that were most predictive of overall similarity judgments on held out memory pairs. In addition to these predictive analyses, descriptive statistics that quantified the correlation between each distance feature and overall similarity judgments were used as another way to identify important features (not illustrated here). See Methods for additional details.

Which dimensions are most important for determining the similarity of human autobiographical memories? As expected, we found that the similarity structure depended on multiple dimensions. However, the emotional characteristics of real-life memories were the most important and reliable organizing feature that we observed. The prominent effect of emotion was found both under conditions of explicit memory search and for more spontaneous recollection. The effects replicated in a separate data set, and they remained strong in an additional experiment which was designed to elicit prosaic, everyday memories from the previous week of the participant’s life. In addition to the effect of emotion, we observed that memories involving similar activities were consistently judged to be similar overall. Finally, to a lesser extent, we found that having a similar spatial environment also made two life memories more similar.

## Results

### Descriptive Features of Real-Life Memories

We asked 513 participants to recall and annotate two memories from their lives. Participants were assigned to one of three conditions: Similar (*N* = 179), Dissimilar (*N* = 164), or Free Choice (*N* = 170). In all three conditions, participants were first asked to recall and describe a specific, memorable event from the last week of their lives (“Memory A”, Figure 2a). Subsequently, participants were asked to recall and describe a second memorable event from any time in their lives (“Memory B”). In the Similar condition, participants were asked for a second memory that was *similar* to the first. In the Dissimilar condition, they were asked for a second memory that was *dissimilar* to the first. In the Free Choice condition, they were asked for any second memory, with no instruction about its relationship to the first. After providing the two memories, each participant provided a series of annotations of their own memories, indicating their spatial, social, temporal, purposive, and emotional properties. The wording of the memory elicitation prompts is shown in Figure 2a, example annotation prompts are shown in Table 1, and the full set is provided in Table S1^1^.

**Table 1:**
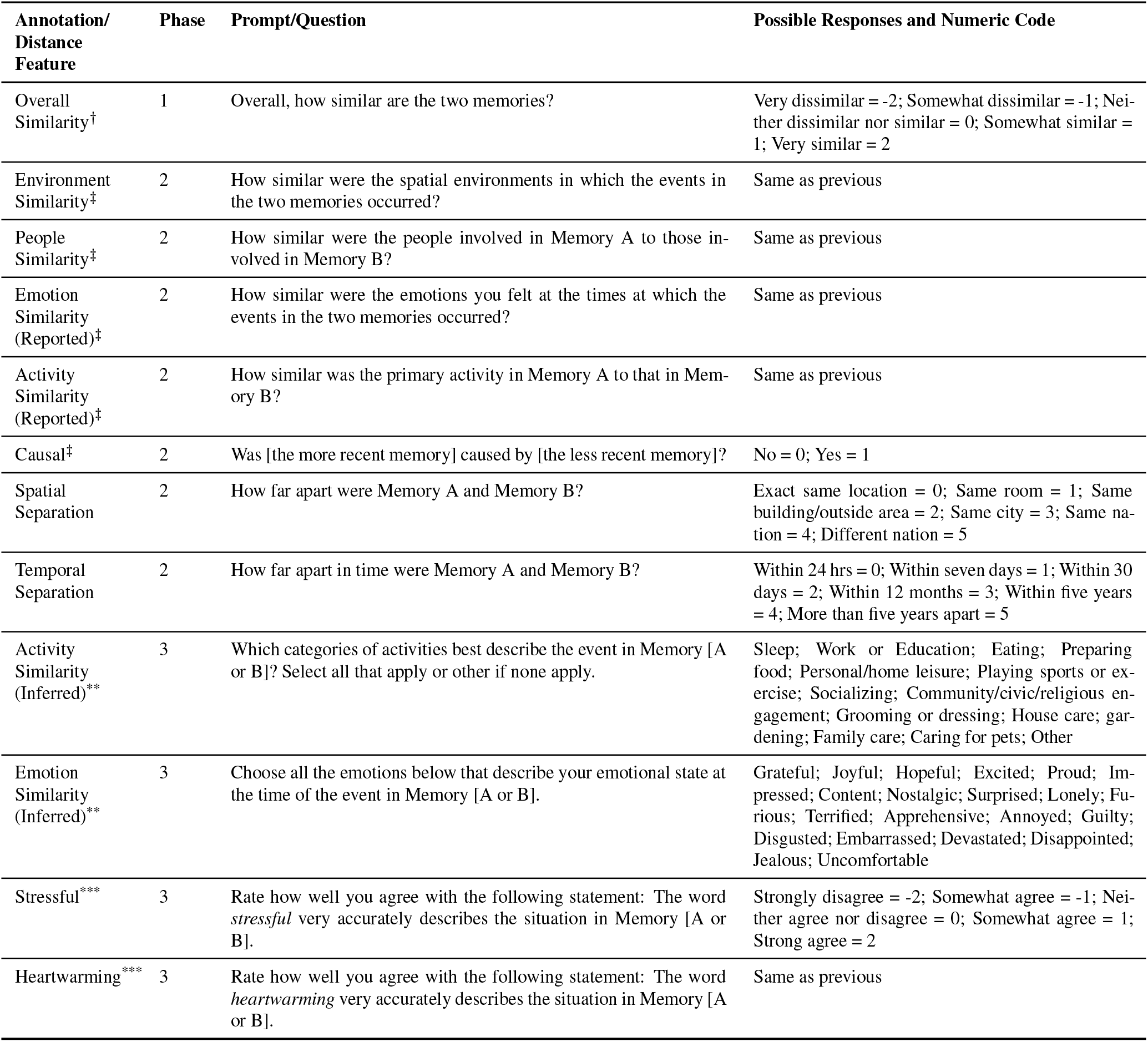
A selected subset of distance features derived from memory annotations. Legend: ^*†*^Overall Similarity is not a distance feature, but the response variable that was predicted by or correlated with distance features in the Free Choice condition. ^*‡*^Indicates this distance feature was computed as the negative of the annotation response. ^**^Indicates this distance feature was computed as the Jaccard distance between the annotation responses for Memory A and Memory B. ^***^Indicates this distance feature was computed as the Euclidean distance between the annotation responses for Memory A and Memory B. Annotations with no marker were directly taken as distance features.

| Annotation/<br>Distance<br>Feature | Phase | Prompt/Question | Possible Responses and Numeric Code |
| --- | --- | --- | --- |
| Overall Similarity<sup>†</sup> | 1 | Overall, how similar are the two memories? | Very dissimilar = -2; Somewhat dissimilar = -1; Neither dissimilar nor similar = 0; Somewhat similar = 1; Very similar = 2 |
| Environment Similarity<sup>‡</sup> | 2 | How similar were the spatial environments in which the events in the two memories occurred? | Same as previous |
| People Similarity<sup>‡</sup> | 2 | How similar were the people involved in Memory A to those involved in Memory B? | Same as previous |
| Emotion Similarity (Reported)<sup>‡</sup> | 2 | How similar were the emotions you felt at the times at which the events in the two memories occurred? | Same as previous |
| Activity Similarity (Reported)<sup>‡</sup> | 2 | How similar was the primary activity in Memory A to that in Memory B? | Same as previous |
| Causal<sup>‡</sup> | 2 | Was [the more recent memory] caused by [the less recent memory]? | No = 0; Yes = 1 |
| Spatial Separation | 2 | How far apart were Memory A and Memory B? | Exact same location = 0; Same room = 1; Same building/outside area = 2; Same city = 3; Same nation = 4; Different nation = 5 |
| Temporal Separation | 2 | How far apart in time were Memory A and Memory B? | Within 24 hrs = 0; Within seven days = 1; Within 30 days = 2; Within 12 months = 3; Within five years = 4; More than five years apart = 5 |
| Activity Similarity (Inferred)<sup>**</sup> | 3 | Which categories of activities best describe the event in Memory [A or B]? Select all that apply or other if none apply. | Sleep; Work or Education; Eating; Preparing food; Personal/home leisure; Playing sports or exercise; Socializing; Community/civic/religious engagement; Grooming or dressing; House care; gardening; Family care; Caring for pets; Other |
| Emotion Similarity (Inferred)<sup>**</sup> | 3 | Choose all the emotions below that describe your emotional state at the time of the event in Memory [A or B]. | Grateful; Joyful; Hopeful; Excited; Proud; Impressed; Content; Nostalgic; Surprised; Lonely; Furious; Terrified; Apprehensive; Annoyed; Guilty; Disgusted; Embarrassed; Devastated; Disappointed; Jealous; Uncomfortable |
| Stressful<sup>***</sup> | 3 | Rate how well you agree with the following statement: The word *stressful* very accurately describes the situation in Memory [A or B]. | Strongly disagree = -2; Somewhat agree = -1; Neither agree nor disagree = 0; Somewhat agree = 1; Strong agree = 2 |
| Heartwarming<sup>***</sup> | 3 | Rate how well you agree with the following statement: The word *heartwarming* very accurately describes the situation in Memory [A or B]. | Same as previous |

In response to the memory elicitation prompts, participants described real-world events with temporal, spatial, social, and self-relevant detail (see examples in Figure 2a). The individual events that were recalled typically occurred in a single location (71% of memories occurred within a space no larger than a building) and over the course of hours (79% of memories were of events lasting no more than six hours).

Combined across the Similar, Dissimilar, and Free Choice conditions, participants generated descriptions with a median length of 54 words (interquartile range: [42-72] words), and median Flesch-Kincaid reading ease score of 84 (interquartile range: [76-90]) (Figure S2). The median word count and reading ease did not significantly vary either between the first and second memories or between conditions (Two-way ANOVA of Memory Position [A/B] and Recall Condition [Similar/Dissimilar/Free Choice], Memory Position: *F* = 1.75, *p* = 0.19; Recall Condition: *F* = 1.05, *p* = 0.35; Memory Position * Recall Condition: *F* = 2.14, *p* = 0.12).

### Which Memory Features Best Predict Overall Memory Similarity Judgments?

Given the large and diverse set of annotations describing participants’ pairs of memories, we sought to determine if any single memory feature could reliably distinguish memory pairs generated in the Similar versus Dissimilar conditions. We reasoned that if an aspect or dimension of an autobiographical memory is important for determining its similarity relationships, then two memories generated in the Similar condition should be closer to each other along that dimension relative to two memories generated in the Dissimilar condition. For example, if the similarity of two memories depends on their relative spatial location, then there should be a tendency for participants in the Similar condition to provide memories that are closer (with smaller ratings for spatial distance) and participants in the Dissimilar condition to provide memories that are further from one another (with larger ratings for spatial distance).

To measure the closeness of memories along specific dimensions, we derived “distance features” from the annotation responses, with smaller observed values of a distance feature indicating being closer along a specific dimension. For some annotations (e.g. “Environment Similarity”) participants directly provided an estimate of the similarity of two memories, which was converted to a distance measure. For other annotations (e.g. “Stressful”) the rating was provided separately for each memory, and so we computed the difference between those ratings. Thus, the “Environment Similarity” distance feature represents the negative of the “Environment Similarity” annotation (Table 1), while the “Stressful” distance feature represents the Euclidean distance between the Memory A and Memory B ratings of the “Stressful” annotation (Table 1). Details for how every distance feature was derived can be found in the Methods section.

We identified important memory dimensions using two approaches: (1) a descriptive statistics approach, which quantifies the tendency for a distance feature to be smaller for Similar memory pairs relative to Dissimilar memory pairs and (2) a univariate predictive (classification) approach, which quantifies how accurately a single distance feature can predict whether a held-out memory pair was generated in the Similar or Dissimilar condition. The intuition for the classification approach is that a distance feature that is important for making similarity judgments will consistently be larger in value for Dissimilar memory pairs compared to Similar ones, and thus, will be diagnostic of which condition a memory pair was generated from. A key benefit of the classification approach is that it tests the generalizability of diagnostic features to unseen data.

For the descriptive statistics, we employed Cliff’s *δ* as a measure of how often each distance feature was smaller for memory pairs from the Similar condition compared to the Dissimilar condition. A Cliff’s *δ* of −1 for a distance feature indicates that distance feature values for all Similar memory pairs are greater than those for all Dissimilar memory pairs, while a Cliff’s *δ* of +1 indicates that distance feature values for all Dissimilar memory pairs are greater than those for all Similar memory pairs. Thus, important features should have Cliff’s *δ* at least greater than zero, with the most important ones being closest to +1. For the classification approach, we employed 10-fold cross-validation classification accuracy as a measure of how well each distance feature predicts whether a held-out memory pair was generated in the Similar or Dissimilar condition. The binary classification was performed using a random forest classifier (see Methods). Overall, the descriptive statistics and the classifier approach produced similar findings (see below) regarding which memory dimensions were most strongly associated with the overall similarity.

Emotion Similarity (Reported) was the strongest indicator of whether a memory pair was produced in the Similar or Dissimilar condition [Cliff’s *δ* = 0.85; *p* ≪ 0.001] (Figure 3a,c; Figure S3a). Concretely, 88% of participants in the Dissimilar condition rated their memory pairs to be somewhat or very emotionally dissimilar, while 81% of participants in the Similar condition rated their memory pairs to be somewhat or very emotionally similar. As a result, the single dimension of Emotion Similarity was sufficient to classify held-out memory pairs (Similar vs. Dissimilar condition) with 85% cross-validated accuracy [IQR 82-91%]. Another measure of shared emotional content, Emotion Similarity (Inferred), was also found to be highly diagnostic of experimental condition. While the Emotion Similarity (Reported) distance feature was derived from participants direct ratings of emotional similarity on a five-point scale, the Emotion Similarity (Inferred) distance feature reflects the number of participant-assigned emotional labels that differ across the two memories. The Emotion Similarity (Inferred) distance feature tended to be greater for memory pairs generated from the Dissimilar condition compared to the Similar condition [Cliff’s *δ* = 0.73; *p* ≪ 0.001], achieving 81% [IQR 76-85%] classification accuracy.

**Figure 3:**
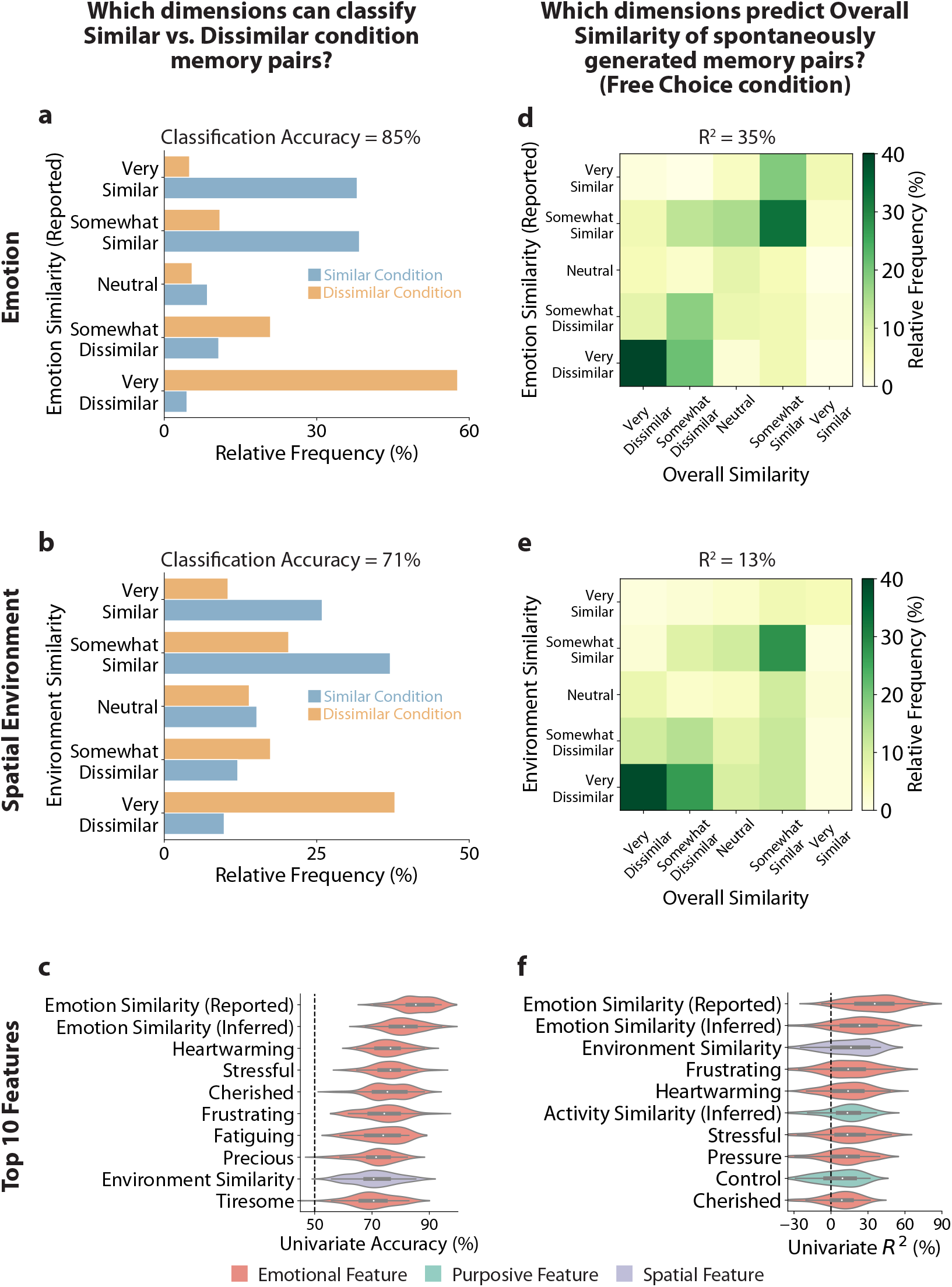
Emotion similarity was the best predictor of memory similarity. **(a, b):** Distributions of ratings of (a) Emotion Similarity (Reported) and (b) Environment Similarity for the Similar and Dissimilar conditions. The separation of the distributions across conditions supports classification with 86% accuracy on held-out samples using Emotion Similarity (Reported) alone, and 71% accuracy using Environment Similarity alone. **(c):** The top ten features are sorted from best to worst classification accuracy. **(d, e):** Covariance patterns of Overall Similarity annotation ratings with (d) Emotion Similarity (Reported) and (e) Environment Similarity annotation ratings for the Free Choice condition. These data support nonlinear regression capturing 35% of variance (Emotion Similarity) and 13% of variance (Environment Similarity) in held-out data. **(f):** The top ten features are sorted from best to worst regression R^2^. Emotional, activity/purposive, and spatial type features are color-coded red, green, and purple, respectively. Chance predictive performance is indicated by the dashed vertical line.

Spatial Separation and Environment Similarity were not as effective as Emotion Similarity for distinguishing Similar and Dissimilar memories, but they nonetheless provided diagnostic information. When using the Spatial Separation dimension, we could only distinguish Similar memory pairs from Dissimilar ones with 62% classification accuracy [IQR 58% - 66%; Cliff’s *δ* = 0.12; *p* = 0.02]; when using the Environment Similarity dimension, classification accuracy was 71% [IQR 68% - 76%; Cliff’s *δ* = 0.5; *p* ≪ 0.001] (Figure 3b; Figure S3a). Of 1000 bootstrap samples of participants, 100% of the time Emotion Similarity (Reported) had a Cliff’s *δ* greater than those for Spatial Separation and Environment Similarity. For Spatial Separation, in both Similar and Dissimilar conditions participants tended to produce pairs of memories that happened in the same city or same country. However, fewer than 28% of memory pairs were within the same building, even when participants were instructed to recall Similar memories. Overall, the spatial proximity of memories was affected very little by whether participants were asked to generate Similar or Dissimilar memories. Regarding Environment Similarity, a substantial proportion (22%) of Similar memories happened in spatial environments that were somewhat or very dissimilar, while a substantial proportion (31%) of Dissimilar memories happened in spatial environments that were somewhat or very similar.

Of the ten most individually predictive features, nine were closely linked to the emotional features of an event (Figure 3c). For example, participants’ ratings of whether each event was “Heartwarming”, “Stressful” or “Cherished” were the 3rd, 4th, and 5th most individually predictive features, behind the overall Emotion Similarity (Reported) and Emotion Similarity (Inferred). The five most accurate non-emotional features were Environment Similarity, Helpful, Useful, Risk, and Activity Similarity (Inferred).

### Controlling for the Effect of Explicit Search Strategies

Emotion Similarity was highly diagnostic of whether participants generated memories in the Similar or Dissimilar conditions, but might this result have been driven by interpretation of the instructions or by the memory search strategies participants employed? In order to generalize the finding to a more natural transition between sequentially retrieved memories, we analyzed responses by a separate group of participants subject to a different set of instructions. In particular, we analyzed graded similarity ratings made by participants in the Free Choice condition, in which participants simply retrieved a second memory after their first memory, without any instruction to find memories that were similar or dissimilar. If Emotion Similarity is truly a diagnostic feature, then it should predict participants’ own overall memory similarity ratings (“Overall Similarity” annotation), for memory pairs that they spontaneously retrieved. In other words, we expect that the similarity of important memory features (i.e. those most important for memory organization and search) will be more correlated with overall memory similarity.

In analyzing the Free Choice data, we again identified important distance features using both descriptive statistics and a predictive approach (nonlinear regression). For the descriptive statistics approach,

Spearman’s *ρ* was employed as a measure of correlation between each distance feature and Overall Similarity. More important distance features should be those that have a stronger tendency to exhibit small values for memory pairs with high Overall Similarity and large values for memory pairs with low Overall Similarity, thus having a Spearman’s *ρ* closer to −1. However, for convenience, we flip the sign of Spearman’s *ρ* reported in the text and figures. Thus, more important distance features should have Spearman’s *ρ* (after sign-flipping) closer to +1. For the regression approach, out-of-sample *R*^2^ was used as a measure of how well each distance feature predicts Overall Similarity for a held-out memory pair from the Free Choice condition (using a random forest regressor).

Our analysis of the Free Choice memory pairs again revealed Emotion Similarity (Reported) as the feature most predictive of Overall Similarity and most correlated with Overall Similarity [out-of-sample *R*^2^ = 35%; Spearman’s *ρ* = 0.61, *p* ≪ 0.001] (Figure 3d,f; Figure S3b). Environment Similarity was also significantly predictive of and correlated with Overall Similarity [*R*^2^ = 16%; Spearman’s *ρ* = 0.45, *p* ≪ 0.001] (Figure 3e,f), but less so than Emotion Similarity; of 1000 bootstrap samples of participants, 99.5% of the time Emotion Similarity had a Spearman’s *ρ* greater than that of Environment Similarity. Spatial Separation, on the other hand, was far less important [*R*^2^ = 2%; Spearman’s *ρ* = 0.16, *p* = 0.008]. Furthermore, consistent with the univariate classification results, the regression analyses revealed that emotional features were common amongst the top features (Figure 3f): of the ten most individually predictive and correlated features, seven were emotional (e.g., Emotion Similarity, Stressful, Heartwarming). Purposive features, specifically Activity Similarity (Inferred) and Control, and Environment Similarity were also important predictors. Together, the univariate classification and regression analyses indicate that emotional states are the most significant factor determining the overall similarity of autobiographical memories, regardless of whether participants are performing directed memory search or an undirected retrieval.

While we observed a reliable contribution of Environment Similarity, the similarity of two memories was not significantly modulated by their exact spatiotemporal coordinates (Spatial Separation and Temporal Separation: out-of-sample *R*^2^ values were 2% and 1%, respectively; Figure S8). Thus, the similarity of two events that occur at the same place appears to derive from the location as a context (with associated sensory and semantic cues), rather than from an absolute coordinate system in space and time.

We did not find a strong influence of whether one event caused the other: only 7% of generated memory pairs in the Free Choice condition were causally linked (determined by a “yes” response to the Causal annotation; Table 1). Among the 12 Free Choice memory pairs that were causally linked, only seven of them were rated somewhat similar or very similar. Hence, the Causal feature was unable to predict Overall Similarity of Free Choice memory pairs (median *R*^2^ = 1%; Figure S8).

We observed a small but significant contribution to memory similarity from the social overlap of two memories, consistent with person-focused theories of memory [Hastie, 1988, Srull and Wyer, 1989]. The out-of-sample *R*^2^ value for the Shared People feature was 7% (Figure S8). In future work, it may be possible to strengthen this prediction by taking into account the specific social relationship at play (e.g. kinship) and the nature of the social interaction.

### Identifying Combinations of Features That Predict Memory Similarity Judgments

Because realworld memories are complex, multiple features will contribute and interact to generate a judgment of overall similarity. Therefore, we designed multivariate analyses to investigate (i) which combinations of features are most effective in classifying and predicting memory similarity; and (ii) how much information is gained by considering multiple features, relative to individual memory features. In addition, multivariate analysis can reveal the extent to which different features carry shared or unique predictive information.

Applying a multivariate model selection procedure, we identified a “top classifier” that employed nine features in a random forest model (see Methods). The contribution of each feature to the multivariate random forest classifier was estimated through permutation importance (Figure 4a and Methods). Features were then ranked from first to last according to permutation importance.

**Figure 4:**
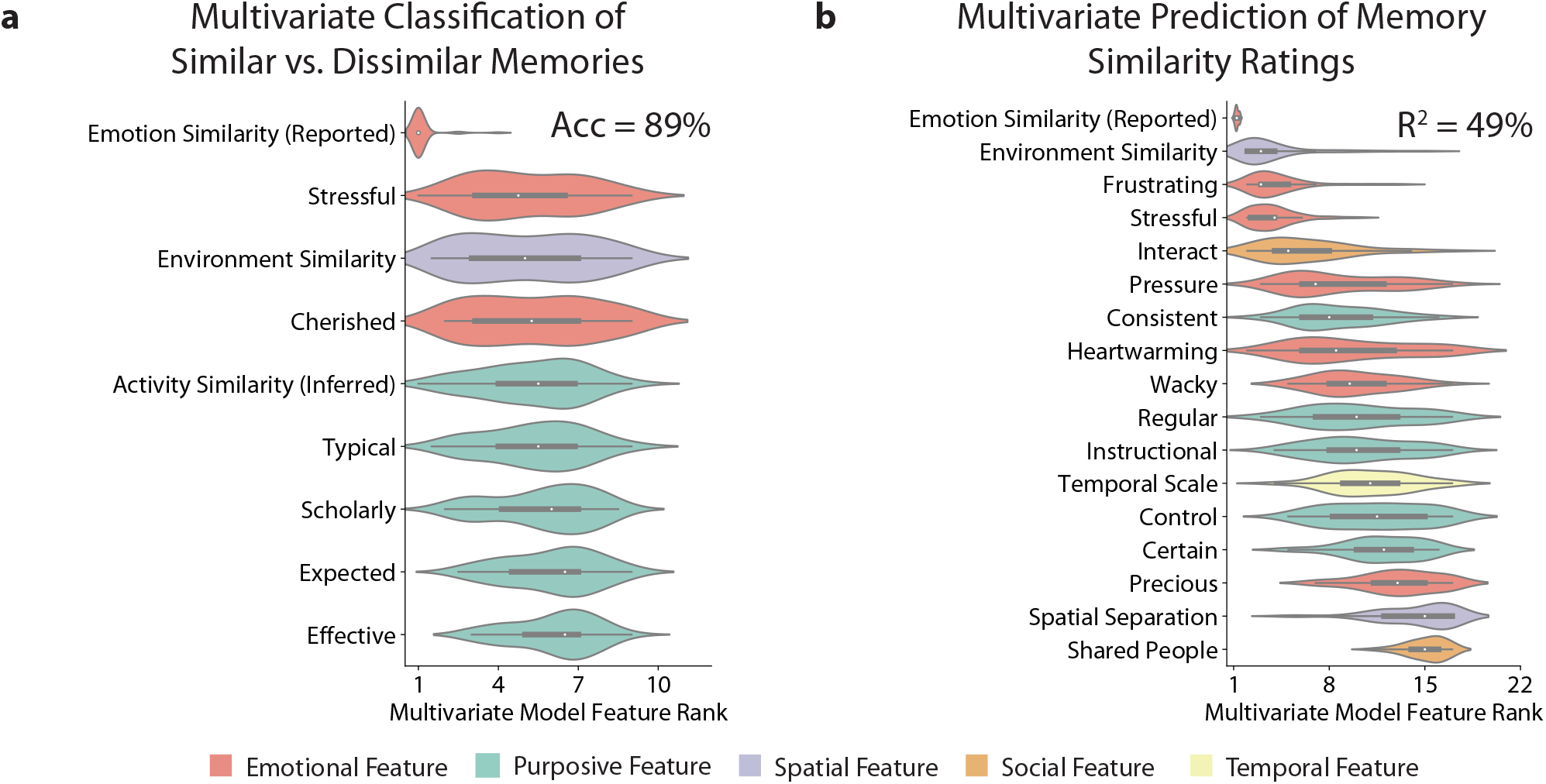
Multivariate predictive models of memory similarity rely most heavily on emotional features. All features were jointly used to train multivariate classification (predicting Similar or Dissimilar condition) and regression (predicting Overall Similarity rating) models. Features were ranked according to estimates of feature importance obtained through the permutation importance method (See Methods). **(a):** Distributions of the feature importance ranks for the subset of features selected to be included in the multivariate Similar vs. Dissimilar classification model. **(b):** The same as **(a)**, but for the Free Choice regression analysis. Multiple features interact to produce better predictive models than the best univariate models, as seen by comparing the multivariate accuracy and R^2^ (89% and 49%, respectively) to those of Emotion Similarity (Reported) alone (85% and 35%, respectively).

Emotion Similarity (Reported) was the most important feature in the multivariate classifier: its median rank was 1, while the next best feature had a median rank of 5. Interestingly, whereas Environment Similarity and Activity Similarity (Inferred) were the ninth and 15th most predictive univariate classifiers, these two features are the third and fifth most important in the multivariate model, respectively; this suggests that the Environment Similarity and Activity Similarity (Inferred) features contain complementary information to the information contained in Emotion Similarity (Reported). Consistent with this interpretation, we note that the multivariate random forest classifier (provided with the nine selected features as input) achieved 89% [IQR 83-94%] accuracy in classifying Similar and Dissimilar memories, and that this was 4% greater than that of the univariate Emotion Similarity (Reported) classifier. Thus, Emotion Similarity (Reported) is the most diagnostic single feature, and it is weakly complemented by other features, for the purposes of predicting overall memory similarity.

Is emotional information necessary for the observed 89% accuracy in distinguishing Similar and Dissimilar memories? We found that non-emotional features could be used to make good predictions of overall similarity, but no combination of non-emotional features was as diagnostic as Emotion Similarity (Reported) alone. When the multivariate classification analysis was repeated with Emotion Similarity (Reported) excluded, and additionally with *all* emotional features excluded, classification accuracy dropped to 85% and 82%, respectively (Figure S4). These emotion-blinded classifiers relied more evenly on all of their selected features; this is unlike the classifier that included Emotion Similarity (Reported), which relied almost entirely on this feature alone.

Finally, we implemented a multivariate regression analysis to predict the overall memory similarity ratings that participants generated in the Free Choice condition. Again applying the model selection procedure, we derived a regression model with 17 features. The out-of-sample R^2^ was 49% [IQR 37-58%]: this performance is 13% greater than the univariate Emotion Similarity (Reported) regressor, confirming that Emotion Similarity (Reported) is complemented by other features in determining graded overall similarity. Three of the features selected were shared with those selected in the previous classification analysis. Emotion Similarity (Reported) was still the most dominant feature (Figure 4b), again having a median rank of 1. The second highest ranked feature was Environment Similarity, which had a median rank of 3.

### Comparison of Everyday Memories and Singular Life Events

Might the strong influence of emotional information on overall memory similarity arise because participants retrieved life memories that were unusual or highly emotional [Christianson and Loftus, 1991]? Since emotionality increases the persistence of memories [Kleinsmith and Kaplan, 1963, Yonelinas and Ritchey, 2015], emotionally powerful memories may be more readily retrievable. If so, then the prominence of emotional information in our analyses could reflect a biased sampling of unusually emotionally rich memories. Of course, if such a selection or attentional effect were occurring, it would still reflect a natural property of real-world memory function – we are studying the memories that people freely retrieved. But in order to understand the role of emotional dimensions in the similarity structure of memories in general, it is important to examine the contribution of emotional dimensions within a set of memories that are not especially emotionally intense. Therefore, we designed a variant of our Free-Choice experiment in which participants generated more prosaic life events, sampled from a randomly-assigned time window on the previous day.

We collected the more prosaic memories using a “Temporally-constrained Free Choice” task. In this task, participants first retrieved a memory from a randomly-assigned two-hour time window from the previous day of their lives (see Methods). The second memory was elicited in the same way as in the original Free Choice condition, and was not temporally constrained. We repeated the multivariate regression analysis on these memory pairs (dubbed “TC-A” for “Temporally Constrained - Memory A”), and we additionally analyzed the subset of these TC-A pairs for which the second memory also occurred within the preceding week (dubbed “TC-AB” for “Temporally Constrained - Memories A+B”)^2^. Participants’ ratings of how well the word “usual” described memory A in the TC-A data were significantly higher compared to the original Free Choice experiment [Cliff’s *δ* = 0.47, *p* ≪ 0.001]. Participants’ ratings of how well “usual” described memory B were significantly higher only for TC-AB [Cliff’s *δ* = 0.55, *p* ≪ 0.001] (Figure S5a,b), consistent with our assumption that temporally-constrained memories would be more prosaic. Additionally, the overall emotional richness of memory A was reduced for TC-A [Cliff’s *δ* = −0.43, *p* ≪ 0.001], while emotional richness of memory B was reduced only for TC-AB [Cliff’s *δ* = −0.47, *p* ≪ 0.001] (Figure S5c,d).

Emotional information remained a strong predictor of memory similarity, even within a new set of more prosaic, everyday memories. In the TC-A analysis, Emotion Similarity (Reported) was the top-ranked feature (Figure S6a); in the TC-AB analysis, the emotional label Cherished was second, and Emotion Similarity (Reported) was third-ranked (Figure S6b), while the top-ranked feature was a marker of whether others were Aware of the events in the memory. Note that the precision of these feature rankings is reduced compared to prior analyses, because these subsets of the data are smaller than the full data set; nonetheless we find that emotional information dominates people’s overall memory similarity judgments, even for these prosaic memories.

### Replication and Extension Data Set

In order to confirm the previous findings, while extending our set of memory features, we collected a separate replication data set of the Free Choice condition (*N* = 99; “Data Set 2”). The first reason for this replication test was that, because we are testing a large number of inter-related memory features, the exact ranks are difficult to measure with high precision. For example, although Emotion Similarity (Reported) was the most predictive single feature, it ranked outside of the top third of predictors for 10% of our cross-validation folds (Figure S7a), and most of of the 66 predictors showed similar levels of variability. Thus, it was important to replicate the basic observation that Emotion Similarity (Reported) was a highly predictive individual feature. The second reason for the replication test was that our initial data collection (i.e. the data described in the previous sections; “Data Set 1”) did not contain explicit probes of the participant-rated people and activity similarities. In particular, the Activity Similarity (Inferred) annotation collected in Data Set 1 was computed based on the number of overlapping action categories in each memory. In contrast, for some other dimensions, such as Emotion Similarity (Reported) and Environment Similarity, we had explicitly asked participants to report similarity estimates along these dimensions. Additionally, Data Set 1 only had one crude measure of the social similarity, which measured the number of people shared across both memories (“Shared People” annotation). To provide a more fair comparison across dimensions, we therefore acquired explicit memory similarity annotations for People Similarity and Activity Similarity (Table 1). The prompts and response style of these new questions were structured in the same way as the questions about Emotion Similarity (Reported) and Environment Similarity.

The findings of our univariate analysis on Data Set 1 (Figure S7a) were broadly replicated in Data Set 2 (Figure S7b). Repeating the univariate regression analysis on the replication data, we found that Emotion Similarity (Reported) remained the feature with highest median feature rank (Figure S7). Additionally, Environment Similarity, Frustrating, Cherished, and Precious were shared in the top-10 predictors across data sets. The Activity Similarity (Inferred) predictor was a top predictor in the first data set, but was replaced by the more explicitly probed Activity Similarity (Reported) measure in the second data set. In fact, Activity Similarity (Reported) was the second-strongest predictor in the second data set, and its predictive performance was comparable to Emotion Similarity (Figure S7b).

### Combining Emotional and Activity Information To Predict Memory Similarity

Activity-related and emotion-related information provide a compact two-variable summary for predicting the similarity of two life events. In a multivariate analysis of the replication data set, we found that 17 features in total were included in the multivariate regression model, with a median cross-validated *R*^2^ of 65%. Emotion Similarity (Reported) and Activity Similarity (Reported) contributed far more than any other features within this multivariate predictive model (Figure 5a). Based on the strong predictivity of these two features, we tested a nonlinear regression model that used only Emotion Similarity and Activity Similarity to jointly predict memory similarity: these two features were strong predictors of Overall Similarity [median out-of-sample *R*^2^ = 56%]. The effectiveness of this simple two-variable model is apparent in a scatter plot of Emotion Similarity (Reported), Activity Similarity (Reported), and Overall Similarity (Figure 5b).

**Figure 5:**
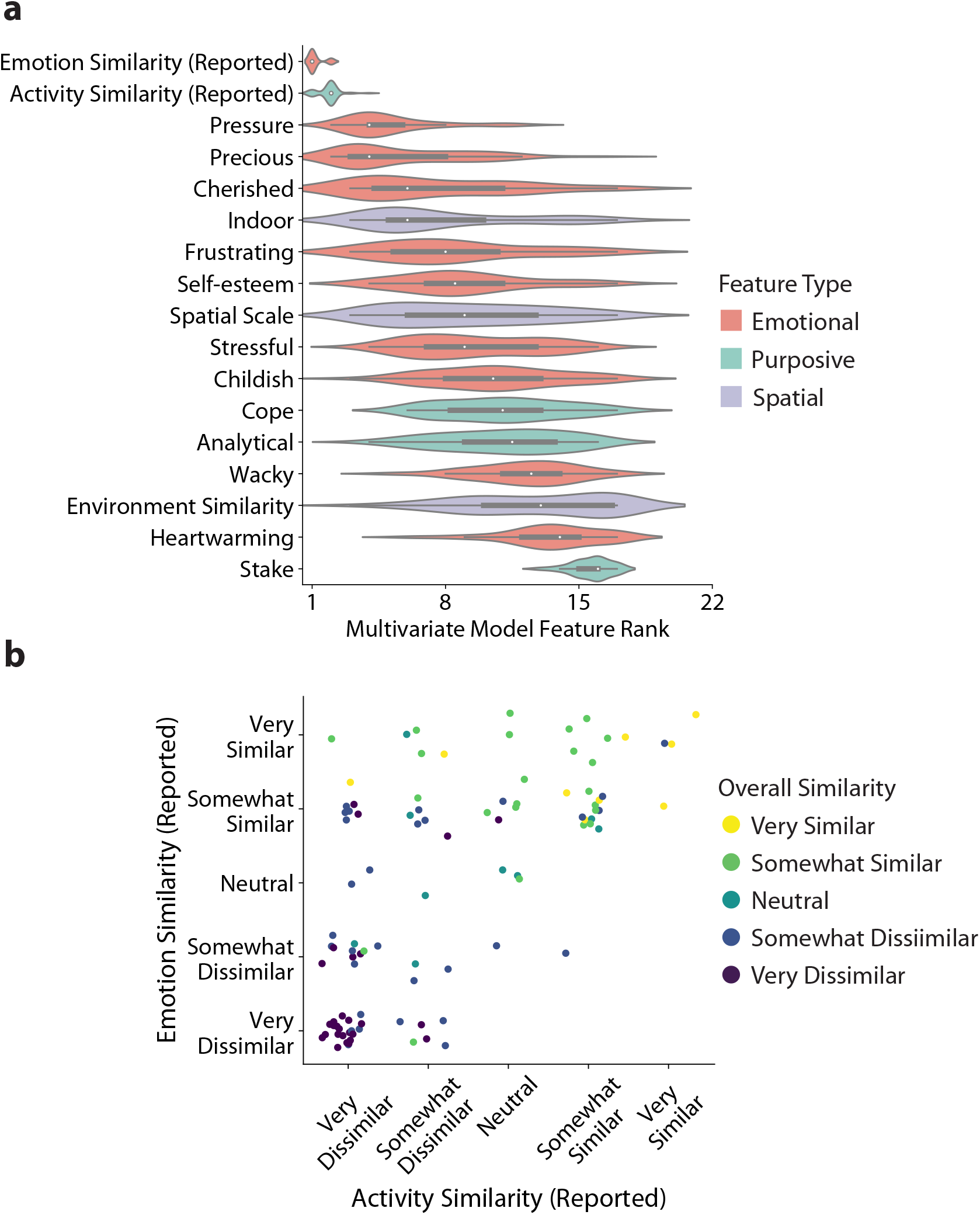
Activity Similarity (Reported) and Emotion Similarity (Reported) are jointly predictive of overall memory pair similarity. **(a):** The same as Figure 4b, but for Data Set 2. **(b):** Scatter plot showing the dependencies between Emotion Similarity (Reported), Activity Similarity (Reported), and Overall Similarity.

## Discussion

How do we mentally organize the memories of our lives? The similarity structure of memories reveals the dimensions along which our life memories will associate, interfere and blend together, and constrains how we search the space of our past experiences. Here we used a data-driven approach to investigate the similarity structure of human autobiographical memory. We found that spatial, temporal, activity-related/purposive, social, and emotional characteristics all contributed to overall memory similarity. However, emotional characteristics were the most consistent single predictor of memory similarity. We replicated the basic finding that emotional features shape our memory organization, and showed that it extended even to prosaic everyday life events. Concretely, this means that people will often understand two life events as similar, even when they occur in different settings, or involve different agents; but people will rarely treat two life events as similar if their emotional content differs.

In addition to the role of emotion, we found purposive features of memory – particularly similarity in activity – to be important predictors of perceived overall similarity of memories. When Activity Similarity was directly elicited from participants (in Data Set 2), it had comparable predictive power to Emotion Similarity (Figure 5). Even when Activity Similarity was only inferred from overlap in Activity labels (Activity Similarity (Inferred) in Data Set 1) it contributed strongly to multivariate prediction (Figure 4). Given the definition of activity as “a sequence of actions performed to achieve a goal,” [Reiser et al., 1985] our findings are consistent with the notion that guidance of future behavior and decision-making is one of three core functions of autobiographical memory [Bluck, 2003, Bluck et al., 2005]. Moreover, our findings also support “script” and “schema” theories in which an overall action or activity, such as “going on a date,” is the central feature of a life memory [Kolodner, 1983, Reiser et al., 1985, Schank, 1983].

Emotional and purposive features of episodic memories may be useful, in combination, for event-level decision making and planning. A bivariate model including the Emotion Similarity (Reported) and Activity Similarity (Reported) features predicted 56% of the variation in the perceived overall similarity of held-out memory pairs. Moreover, participants in the Dissimilar condition explicitly reported emotion as the most common dimension along which their two memories were dissimilar, and participants in the Similar condition explicitly reported emotion and activity as dimensions along which their two memories were similar more frequently than they mentioned social, spatial, temporal, or causal dimensions (Figure S9). Consider that we often face situations similar in activity and affective experience to ones we have encountered in the past, and therefore look to past experiences to help us navigate new ones. For example, a person preparing for a first date might look back to previous first-dates and reflect on why they went well or badly. While prior experiences with similar locations and times may provide information that is relevant and useful, perhaps the more informative features of the prior experiences relate to emotions (“when did I last feel this way?”) and to actions and goals (“when did I last try to impress someone with my cooking?”).

Consistent with theories that emphasize spatial scaffolding of memory representations [Robin et al., 2016], we also found that the spatial context of a life event was an important organizing factor for searching autobiographical memory: Environment Similarity reliably ranked in the top 10 of the 66 tested features (Figure 3). In contrast, we found very small effects of the absolute spatiotemporal coordinates of the events. Therefore, the effects of spatial context appear to derive from properties or cues associated with the locations (e.g. two memories are similar because they both occurred in a kitchen).

Emotion Similarity (Reported) was the single most consistent predictor of Overall Similarity, and this effect persisted even for less emotional and more prosaic life memories (Figure S6). Earlier mapping of the similarity structure of human autobiographical memory [Brown and Schopflocher, 1998] explicitly excluded affective annotations of participants’ memories. Later work argued that personal relevance and emotional valence and arousal could be important for the clustering of life memories [Wright and Nunn, 2000], and that the valence of autobiographical memories affects how they are retrieved [Sheldon and Donahue, 2017]. The present data enabled us to demonstrate the importance of emotional properties of memories in a quantitative manner, comparing emotion against other theoretically prominent dimensions, such as the spatial, temporal and social features of memories.

The affective valence of memories made a large contribution to their overall memory similarity. Similarity in emotional content remained a strong predictor, even when it was elicited only from endorsement of a single emotional adjective (e.g. whether both memories were similarly “Heartwarming” or “Stressful”; Figures 3c,f, S3). The predictive power of individual affective labels is consistent with the finding that “valence” and “adversity” are amongst the most important dimensions for psychologically summarizing situations [Parrigon et al., 2017, Rauthmann et al., 2014]. Examining the specific emotional labels assigned to Memory A and Memory B (Figure S10), it appears that participants in the Similar condition produced memories of the same valence while participants in the Dissimilar condition produced memories of opposite valence, as reflected in the block structure in Figures S10a,b.

However, it is not only the emotional intensity or valence of memories that matter to their later recollection, but also their specific emotional qualities. It is established that more emotional events produce more lasting memories [Hamann, 2001, Talarico et al., 2004], an effect which likely depends on interactions between the hippocampus and amygdala. Moreover, we know that retrieval is modulated by the emotional state at the time of retrieval [Bowen et al., 2018, Holland and Kensinger, 2010, LaBar and Cabeza, 2006, Palombo et al., 2016, Phelps, 2004, Talmi, 2013]. Memories with exactly matching emotion labels were more likely to be judged as similar and to be retrieved in relation to one another (see Emotion Similarity (Inferred) in Figure 3c,f, a measure of overlapping emotion labels), and this effect was not simply due to them having the same valence (note the diagonal structure in Figure S10a). Therefore, our life memories are not only “tagged” as emotional or unemotional, but are associated with (and may be later retrieved via) specific emotional features. Altogether, the data are consistent with the idea that rich emotion representations influence memory and decision-making in real life-settings, and that emotion must be incorporated into computational models of these processes [FeldmanHall and Chang, 2018, Huys et al., 2015, Murty et al., 2020, Talmi et al., 2019].

Given the high-dimensionality of episodic memories [Cowell et al., 2019], a dominant contribution from a single dimension, like emotion, is somewhat surprising. One possible explanation is that emotional appraisals provide compact, lower-dimensional representations of situational actions and outcomes [Arnold, 1960, Lazarus and Lazarus, 1994, Man and Cunningham, 2019, Solomon, 1993]. The neural and cognitive process of emotional appraisal involves abstract situational features that cannot be readily derived from simpler characteristics such as facial features or semantic-valence associations [Barrett, 2017, Satpute and Lindquist, 2019, Skerry and Saxe, 2015]. Consistent with the notion that emotions provide a compact event summary, we found that no multivariate combination of multiple non-emotional features was as diagnostic as the single feature of Emotion Similarity (Figure S4). The data are consistent with a key claim from appraisal theories of emotion: *“The idea, common to all appraisal approaches, is that there is a deep structure or underlying constancy in situations that makes them sources of anger, fear, or joy”* [Clore and Ortony, 2008]. If such a “deep structure” exists, then it could be advantageous for our memory systems to use such structure to retrieve situationally relevant memories, which could support (explicit) reasoning and (implicit) outcome sampling [Bornstein and Norman, 2017] from related scenarios.

The notion that emotional memory provides a “deep structure” or summary representations for events is consistent with neurocognitive models of memory organization. In the human brain, emotional and motivational processing are thought to involve the anterior hippocampus [Fanselow and Dong, 2010] and the “AT system,” a network of brain regions centered in the anterior-temporal lobe [Ranganath and Ritchey, 2012]. The anterior temporal lobe and anterior hippocampus are also thought to support coarser-grained, integrative representations, in contrast to the more distinctive and idiosyncratic episodic detail whose recollection relies on posterior temporal and posterior hippocampal processes [Brunec et al., 2018, Poppenk et al., 2013]. In the specific context of autobiographical memory, more posterior memory circuits may support a “perceptual” form of memory, focused on detail and vivid reexperiencing, while anterior circuits may support a more “conceptual” aspect of autobiographical memory, useful for decision-making in unstructured and unfamiliar scenarios [Ranganath and Ritchey, 2012, Ritchey and Cooper, 2020, Sheldon et al., 2019]. Similarly, it has been proposed that the anterior hippocampal subregions, closer to the amygdala, have a particular role for the initial, more abstract construction of an autobiographical memory, whereas more posterior regions are involved in the more detailed elaboration of the memory [McCormick et al., 2015, Morton et al., 2017]. Thus, in all of these neurocognitive models, emotional processing, involving more anterior memory networks, is naturally linked to more abstract representations which may correspond to the “deep” structure of events.

Future work will be required to determine whether emotional associations exert an outsize influence only on self-relevant autobiographical memories. Technically, all episodic memories are autobiographical, but memories vary greatly in their relevance to our notions of self and our life narratives. The episodes studied in standard memory paradigms usually have little self-relevance (e.g. encoding a sequence of unrelated words; receiving sugar water) whereas our daily lives contain many events with self-relevance (e.g. searching for a new apartment; having lunch with a friend). Previous work suggests that information encoded in a self-referential manner is remembered better than information encoded non-self-referentially [Durbin et al., 2017], but what influence does the degree of self-referential encoding have on the associative structure of memory? It remains to be seen whether the powerful role of emotional content is specific to highly self-relevant memories or whether the role of emotions would equally extend to complex fictional scenarios (e.g. the events in a recurring television series) which could be as complex as real events, but less personally relevant.

A second point for future work will be to determine how the similarity structure of memory depends on the purpose for which memory is being used and the availability of endogenous or exogenous cues. For example, when presented with a strong exogenous cue (e.g. seeing an apartment building you had lived in decades before), you may automatically retrieve other memories related to that location, and spatial proximity will be the primary factor in that memory search (for the importance of cue familiarity see Robin et al. [2019]). On the other hand, when memory is being effortfully searched and the primary cue is endogenous (e.g. when we are reminiscing about a reunion dinner) then the particular location of the dinner may be less important, relative to the emotional, social and purposive features associated with the dinner. More generally, the particular manner in which memory is used will matter: the absolute spatial position of an event may be much more important in the context of a navigation task; the causal flow and chronological order may be more prominent organizing factors when a person is narrating a long sequence of inter-related life events [Barsalou, 1988, Linton, 1986, Moreton and Ward, 2010, Uitvlugt and Healey, 2019]. In this experiment, we measured the similarity structure in the context of a fairly generic task, by simply asking participants to recall events from their lives without further instruction, and it will be important to cross-reference these results against other elicitation procedures. We hypothesize that spatial context would exert stronger effects in the context of spontaneous and automatic memory cueing, more than in the setting of explicit and effortful memory search.

A final important question for future work is whether our findings will generalize to implicit measures of memory similarity. In the present experiments, we only measured memory similarity directly: participants provided their judgments of memory similarity in response to an explicit question or via an explicit instruction to retrieve similar (or dissimilar) memories. It will be important to test whether the similarity structure we found would replicate when using indirect techniques for measuring memory similarity, for example by measuring memory interference or confusability, or the biases in reaction times used in the study of event clustering [Wright and Nunn, 2000].

In summary, we find that while the objective features of events (“who”, “when,” and “where”) certainly link events in our memories, the most powerful explicit associations depend on “what you were doing” and “how you felt about it”. This work highlights the need for more detailed behavioral and neural measurements of how rich emotional appraisals affect the encoding and retrieval of real-world events. In addition, it appears that we require theoretical models of real-world memory that emphasize the role of emotions and situational goals, explaining how they can serve as sources of summary information for rapid retrieval and evaluation of relevant life events. By understanding how emotional and purposive information are woven into our experience and recollection, we can improve the applicability of theoretical memory models to real-world settings, and will advance our tools for practically preserving and recovering memory function.

## Methods

### Autobiographical Memory Similarity Experiment

We collected and scored real autobiographical memories in two stages. In the first stage, participants recruited via the Amazon Mechanical Turk (AMT) platform provided free-form written descriptions of events from their lives (“memory elicitation”; two life events per participant). In the second stage, each participant provided feature labels for their life events and judged their similarity along a number of dimensions (“memory annotations”). Prior to the start of the experiment, participants were randomly assigned to one of three experimental conditions, which will be described in detail in the following paragraph.

In the memory elicitation stage, participants in all three conditions began by providing a single free-form memory of an event from the past week of their lives. They were then prompted to provide a second memory that was similar to the first memory (Similar condition), dissimilar from the first memory (Dissimilar condition), or not explicitly instructed to be related to the first one in any way (Free Choice condition; see Figure 2a for memory elicitation prompts). Responses were required to be a minimum of 150 characters in length. As expected, participants in the Similar group provided memories that were much more closely related than participants in the Dissimilar group (Figure S1). We observed a large and significant difference in the similarity ratings across groups, with memories in the Similar group rated more similar [Cliff’s *δ* = 0.85, *p* ≪ 0.001].

In the subsequent annotation stage, participants rated diverse properties of their memories, including the recency and temporal scale (“when”); the spatial scale and relative spatial proximity (“where”), the primary activity (“what”), as well as the emotional, social, and purposive content of each life event. Continuously throughout the the annotation phase of the experiment, the full text of the memories provided by the participant, was displayed in the top left of the screen (Memory A) and top right of the screen (Memory B). The annotation stage was divided into four sequential phases. *Phase 1* consisted of two trials. In the first trial, participants were asked to elaborate on, in the form of free response, the similarities and/or differences between their two memories, depending on experimental condition. In the second trial, they rated the overall similarity of their memory pairs on a five-point likert scale. *Phase 2* consisted of seven trials (nine for collection of Data Set 2). Each trial involved directly rating distances or similarities between their memories along a different dimension. For example, in one trial participants were asked to rate (on a discrete ordinal scale) the degree of spatial separation between the events in the two memories. In *Phase 3*, each trial involved annotating a particular dimension of just one of their memories. The same 62 dimensions were annotated for both memories individually, resulting in 128 trials. For example, in one trial participants were asked to rate (on a discrete ordinal scale) how long ago the event in Memory A occurred. A separate trial had the participants rate this same temporal dimension but for Memory B. *Phase 4* consisted of two trials. In one trial participants were asked to provide one to five words that best summarize Memory A. The other trial involved doing the same for Memory B.

The order of trials within each of Phases 2-4 was randomized across participants. The randomization procedure used for Phase 3 ensured that no more than three consecutive trials involved annotating the same memory. Each property rated during the annotation stage, with the exclusion of four, was used to derive a corresponding distance feature for the subsequent predictive (classification and regression) and descriptive statistics analyses. Table S1 delineates the prompts for each annotation trial. Seven “catch trials” were randomly inserted into the annotation stage for the purpose of quality control. Catch trials were questions that could be answered with general common knowledge, but required some human-level reasoning in order to assess that participants were paying attention and to assess whether participants might be bots.

### Participants and Exclusion Criteria

892 participants (median age range 30-34, min age range 18-19, max age range 70-74, 374 females) completed our online memory experiment for payment via the Amazon Mechanical Turk platform. Participants gave informed consent in accordance with the requirements of the Johns Hopkins University Homewood Institutional Review Board.

Participants’ responses were excluded from analysis if the responses did not meet all of three criteria. First, participants must have correctly answered at least six out of seven randomized catch trials (Criteria 1). Second, participants assigned to either the Similar or Dissimilar conditions must have had memory pair overall similarity ratings consistent with their assigned condition (Criteria 2). Specifically, participants in the Similar condition must not have rated “Somewhat Dissimilar,” or “Very Dissimilar,” while participants in the Dissimilar condition must not have rated “Somewhat Similar,” or “Very Similar.” Last, participants were excluded if their textual memory responses described anything other than an autobiographical memory (Criteria 3). Exclusions were made using each criteria sequentially in the order described above. That is, a subsequent exclusion criteria was used on the participant pool remaining after the previous exclusion criteria had been applied. For Data Set 1, for the experiments in which Memory A was from the past week, 513 participants remained after exclusion; for the temporally constrained experiments in which Memory A was from the previous day, 120 participants remained after exclusion. For Data Set 2, 99 participants remained after exclusion. Table S2 provides a more detailed summary of the number of participants excluded, broken down by experimental condition and exclusion criteria.

### Construction of Distance Features

The inputs to the classification and regression algorithms were distance features derived from the participants’ ratings and labels of their memories during the annotation stage. Distance features were generated from the raw annotation responses of participants using custom python code. Each distance feature quantifies how far apart two memories of a generated memory pair lie along a particular dimension. The ratings from Phase 2 of the annotation stage directly measured the similarity/distance of Memories A and B along a particular dimension (e.g. Environment Similarity). Since we wanted distance features to be defined such that smaller values correspond to being closer along a particular dimension while larger values correspond to being farther apart along a particular dimension, the Shared People, Causal, Environment Similarity, Emotion Similarity (Reported), Activity Similarity (Reported), and People Similarity distance features were taken to be the negative of their corresponding annotation ratings. For instance, if a participant rated “1” for the Environment Similarity annotation, then the value of the Environment Similarity distance feature for that participant’s memory pair was taken to be “-1.” The remaining Phase 2 distance features were directly taken to be the corresponding annotation ratings. Each of the annotations in Phase 3 of the experiment was performed separately for Memory A and for Memory B. Therefore, derivation of distance features from Phase 3 annotations required defining a distance measure between corresponding Memory A and B ratings for each dimension. For dimensions which were rated on an ordinal scale, such as Heartwarming, the distance measure was the Euclidean distance (i.e. absolute value of the arithmetic difference) between Memory A and B ratings. For dimensions which were annotated by selecting one or more qualitative response options, such as Activity Similarity (Inferred), the Jaccard distance measure was used. Specifically, if *F_A_* and *F_B_* are the sets of response options selected for Memories A and B, respectively, then the Jaccard similarity index, *S_J_*, is defined as the number of elements that *F_A_* and *F_B_* share divided by the number of unique elements in the union of *F_A_* and *F_B_*. The Jaccard distance is then 1 - *S_J_*.

A few of the annotations listed in Table S1 were not used to generate distance features. The response to the “More Recent” annotation merely determined how the “Causal” annotation question was presented to participants. The “Temporal Location” annotation wasn’t used because this measurement was not collected for all participants (the ‘Temporal Separation” annotation, which was used, captures the same information for the purposes of our analyses). The “Emotion Intensity” annotation was used solely to confirm that memories collected in the “Temporally Constrained Free Choice” experiment were less emotional than memories collected in the original baseline experiment. The “Word Cloud” annotation was not used because it isn’t clear what type of dimension (i.e. Temporal, Spatial, Emotional, etc.) its distance feature would belong to.

Note that, because the distance features derived here have variable scaling properties (e.g. some are only ordinal, while others may have interval scaling), it was important to use analytic methods that are invariant to the scaling. This motivated us to use (ordinal) metrics such as Cliff’s *δ*, and Spearman’s rank correlation, as well as nonparametric nonlinear regression and classification procedures which do not make strong assumptions about the scaling properties of their input.

### Classification/Regression Analysis

Univariate classification analyses (Similar and Dissimilar conditions) and regression analyses (Free Choice condition) were performed using random forests [Breiman, 2001]. These analyses (and subsequent multivariate classification/regression analyses) were performed using the R implementation of random forests publicly available in the SPORF package [Tomita et al., 2020]. For each distance feature, a random forest classifier was trained using the feature values for a subset of memory pairs from the Similar and Dissimilar conditions, along with class labels indicating which condition each memory pair was generated from. Classifiers were evaluated based on accuracy in predicting the class labels (Similar or Dissimilar condition) for a separate set of memory pairs. Similarly, for each distance feature, a random forest regressor was trained using the feature values for a subset of memory pairs from the Free Choice condition, along with the Overall Similarity ratings for each memory pair. Regressors were evaluated based on *R*^2^ of Overall Similarity predictions for a separate set of memory pairs. Training and evaluation were done for each distance feature sequentially, using a standard stratified 10-fold cross-validation (CV) sampling procedure. The 10-fold CV was repeated five times, resulting in 50 total CV replicates. Distributions shown in Figure 3 are distributions of the accuracies and *R*^2^ values of each feature across the 50 replicates.

If multiple features contribute to overall memory similarity judgments in an additive manner, or if different features are highlighted for different participants when making similarity judgments, then a multivariate model should be able to predict memory pair similarity better than the best univariate predictor. For multivariate analyses, we trained random forest classifiers or regressors. The input to a classifier or regressor was an *N × d* distance feature matrix, where *N* was the number of participants pertinent to the classification or regression analysis, and *d* was the total number of distance features. CV was performed in the same way as for the univariate analysis.

Feature importance estimates in the multivariate models were computed using the permutation importance method (Breiman 2001). This method follows the logic that the more important a feature is for predicting experimental condition (classification) or overall similarity rating (regression), the worse prediction performance will become when observed values of this feature are permuted. Thus, after the random forest model had been trained for each of the ten CV iterations, each distance feature was sequentially randomly permuted across participants in the held-out set. Then, for each distance feature that was permuted, the accuracy or R^2^ was recomputed. Importance of each distance feature was measured as the decrease in accuracy or R^2^ when that feature was permuted, normalized by that of the intact (no features permuted) model.

In the multivariate prediction scheme, we employed an iterative feature selection procedure to reduce the dimensionality of predictors. The purpose of this dimensionality reduction was two-fold. The first reason was to alleviate overfitting, thereby improving both generalization performance and estimates of feature importances. The second was to identify a compact set of the key predictors of similarity, if such a compact set should exist. Specifically, initial estimates were obtained by including all distance features in the multivariate models, repeating for each of the 50 CV replicates. Features were ranked from one (most important) to *d* (least important) for each CV replicate. For each feature, the number of times out of the 50 CV replicates that it ranked lower (i.e. better) than the average rank 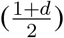 was computed. The 50% of features with the lowest frequency of ranking better than the average rank were removed, and the procedure was repeated with the subset of features. This was performed iteratively four times. Thus, four sets of features, each subsequent set a subset of the previous, were evaluated. The final set of features selected and reported in the results was the one (out of the four nested sets) that achieved the largest median CV accuracy or R^2^. Distributions shown in Figures 4 and 5 are distributions of the ranks of each feature across the 50 replicates.

### Temporally-Constrained Experiment

To elicit memories that were less personally meaningful or emotional, we collected a variant of the Free Choice data set in which the autobiographical memories were constrained to recent time windows. The memory elicitation for the first memory (Memory A) was identical to the of the original Free Choice condition, except that participants were asked to produce a memory from an assigned two-hour time window on the previous day. Each participant was assigned randomly to either the 10am-noon, noon-2pm, 2pm-4pm, or 4pm-6pm time window. The wording of the elicitation for the second memory (Memory B) was identical to that of the original Free Choice condition. Multivariate regression analysis was repeated on this data set (Temporally Constrained Memory A, TC-A). A second regression analysis was performed on the subset of these data for which memory B was no more than a week old (Temporally Constrained Memory A and B, TC-AB). For the purposes of this TC-AB analysis, the recency of Memory B was determined from each participant’s response to the Temporal Location question.

## Footnotes

1 The Activity Similarity (Reported) and People Similarity annotations were not collected for the analysis that follows in this subsection. These annotations were collected in a replication data set, described in a later subsection.

2 Memory B in the TC-AB analysis was not constrained at the time of the experiment. That is, participants could produce a Memory B from any point in their lifetime. However, at the time of conducting the TC-AB analysis, we only examined the cases in which Memory B occurred within the preceding week.

## Data Availability

Textual memory descriptions and responses to the annotation trials can be downloaded from https://github.com/HLab/AutobioMemorySimilarity

## Author Contributions

TMT and CH conceived the study. CH provided statistical input. TMT collected the data and performed the analyses. TMT, MDB, and CH wrote the manuscript. All authors approved the manuscript for peer review.

## Acknowledgments

We are grateful to Nick Diamond, Iva Brunec, Lisa Musz, Buddhika Bellana, Hongmi Lee, and Eduardo Sandoval for helpful feedback and discussions, and to Natalie Wang for preliminary analysis and summarization of participants’ textual memory descriptions. This work was supported by an RCP2 award from the Centre for Aging and Brain Health Innovation (CABHI).

## Supplementary Material

**Figure S1:**
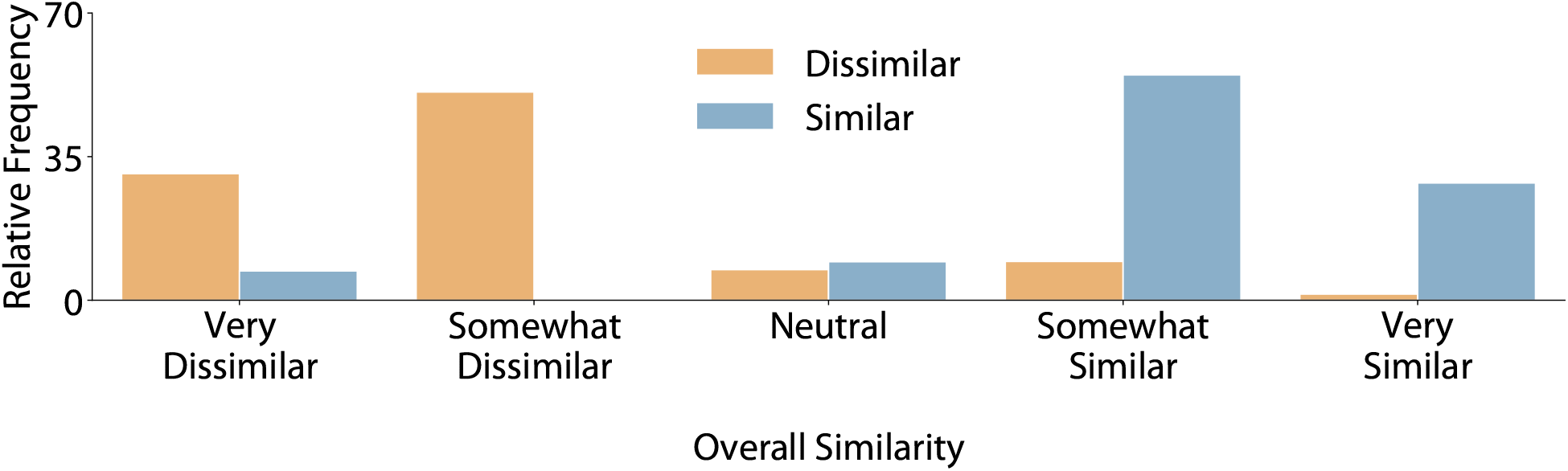
Overall memory pair similarity ratings from participants in the Similar and Dissimilar conditions confirm that their memories were indeed judged to be similar and dissimilar, respectively. Note that the above data were computed from the data of all participants, before exclusion: participants in the Similar condition were excluded from analysis if they reported their memory pairs as being Somewhat or Very Dissimilar, and likewise participants were excluded from the Dissimilar condition if they reported their memories as being Somewhat or Very Similar.

**Figure S2:**
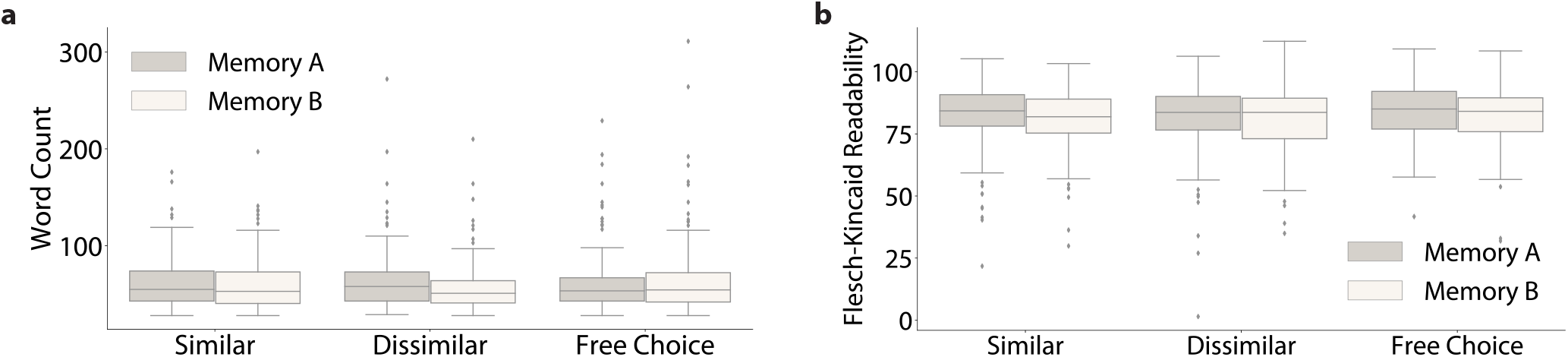
Summary statistics of textual memory descriptions. **(a):** Box plots showing distributions of the numbers of words per memory response. Boxes represent the interquartile range (IQR), the horizontal lines within each box represent the median, whiskers represent 1.5±IQR, and individual points are outliers. **(b):** Box plots showing distributions of the Flesch-Kincaid readability scores of memory responses.

**Figure S3:**
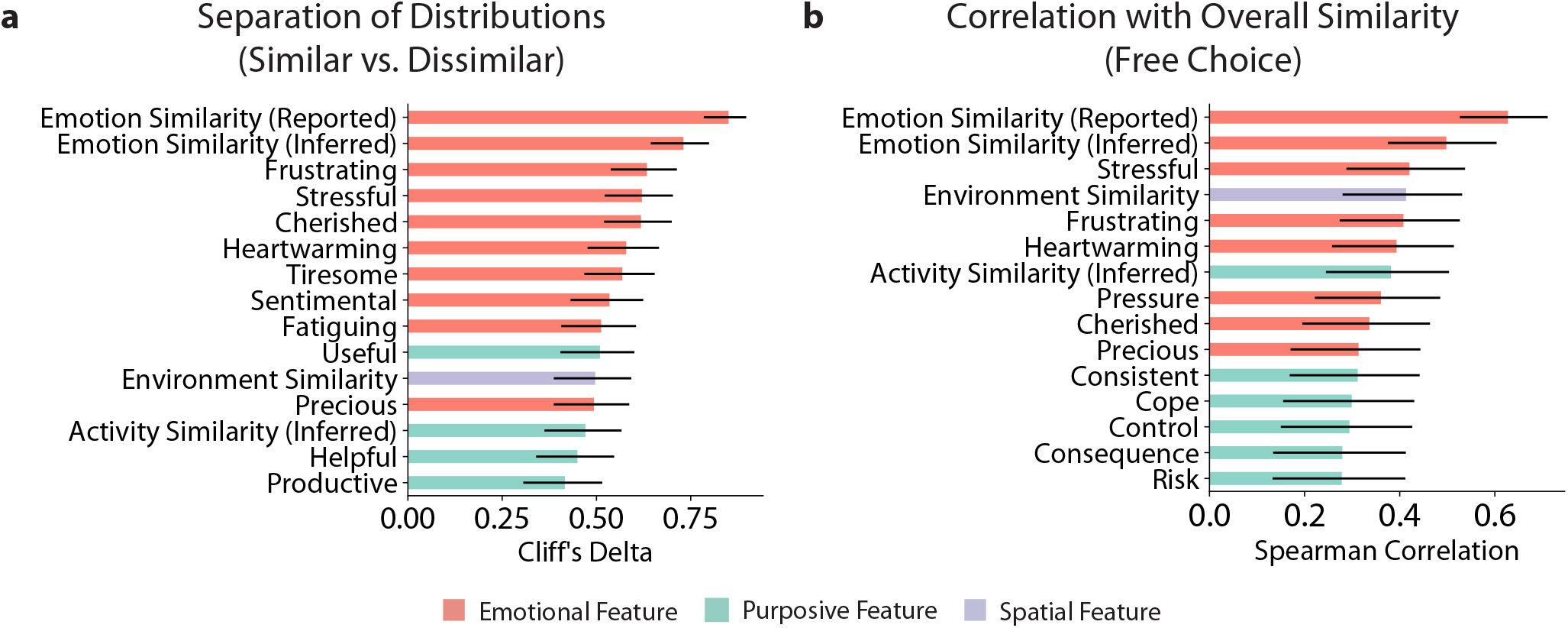
Whole-sample inference using Cliff’s *δ* and Spearman’s *ρ* indicate emotional features are significantly diagnostic of memory similarity. **(a):** The fifteen features with the largest Cliff’s *δ* when comparing the Similar vs. Dissimilar memories, sorted from greatest to least. **(b):** The fifteen distance features that most strongly correlated with overall memory dissimilarity, sorted from greatest Spearman’s *ρ* to least.

**Figure S4:**
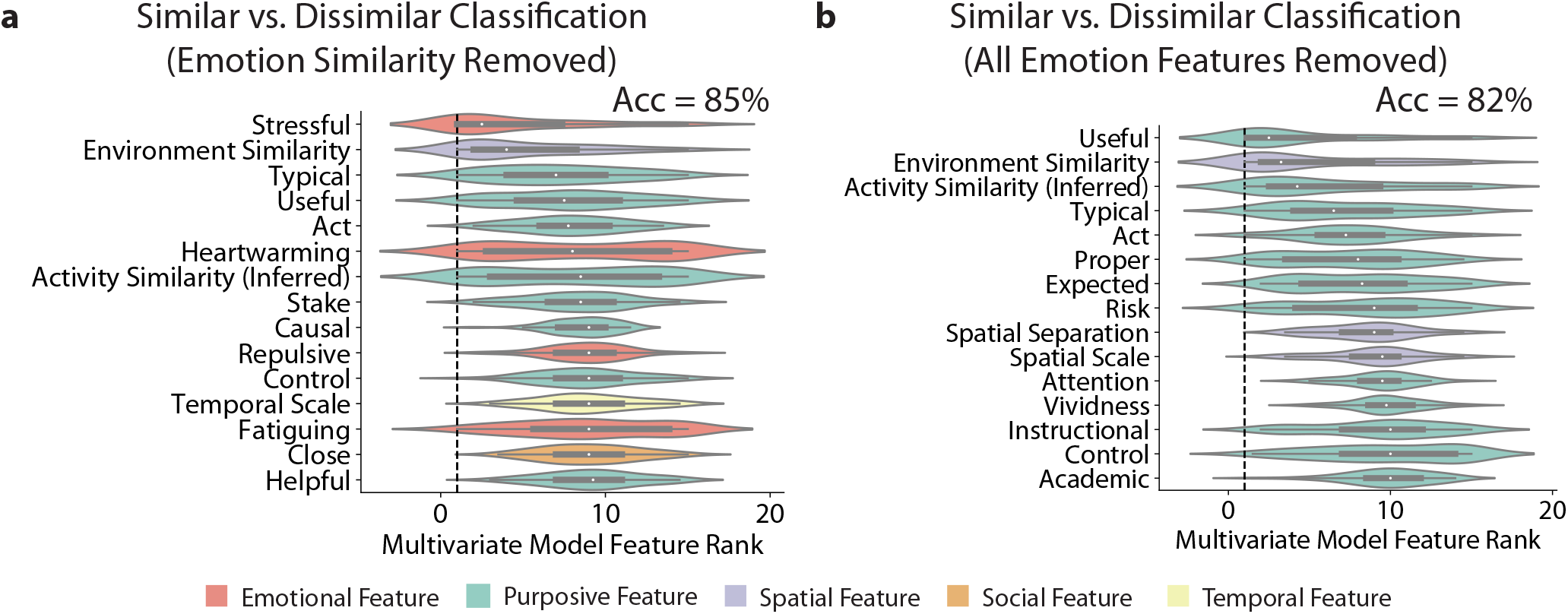
When emotional features are omitted, multivariate classification of Similar and Dissimilar memories is worse, and relies more evenly on multiple features. **(a):** Feature importance ranks of the features selected in the multivariate classification model when Emotion Similarity (Reported) and Emotion Similarity (Inferred) are omitted from analysis. **(b):** Feature importance ranks of the features selected in the multivariate classification model when all emotional features are omitted from analysis.

**Figure S5:**
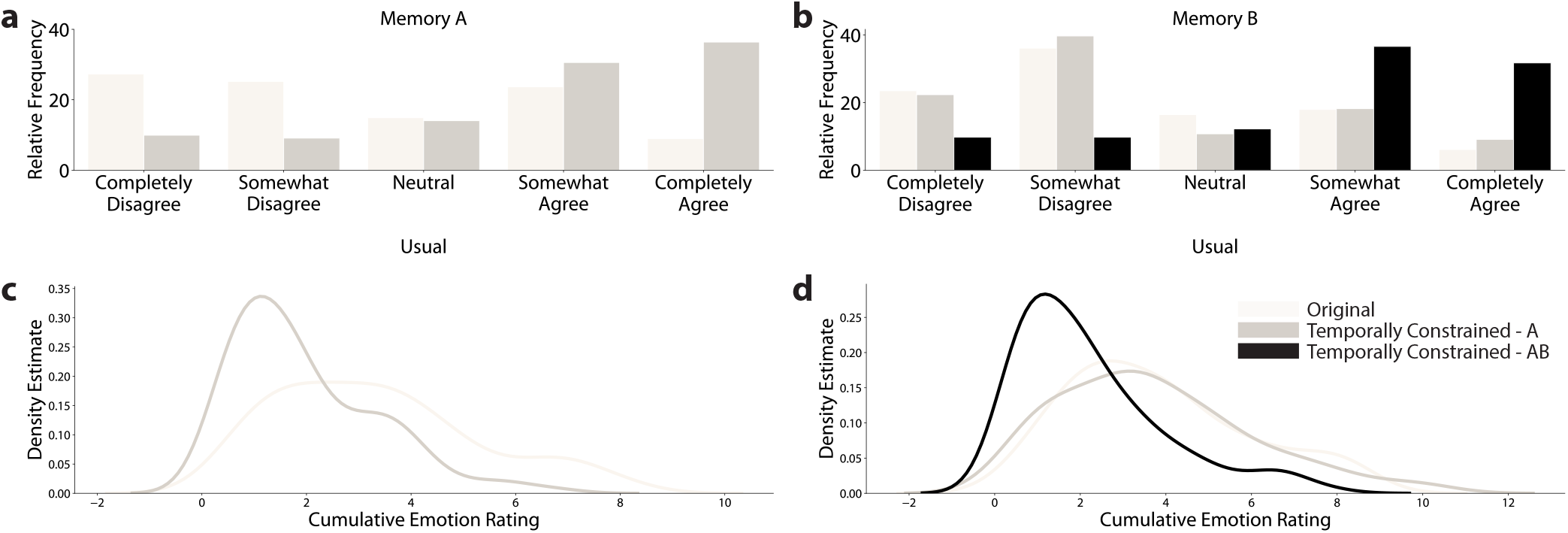
Memories that are temporally constrained to be more recent are rated as more “usual” and less emotional. **(a):** Distribution of ratings for the Usual annotation of Memory A. **(b):** Same as **(a)**, but for Memory B. **(c):** Distribution of the sum of Emotion Intensity for Memory A. **(d):** Same as **(c)** but for Memory B. Distributions are broken down by the original baseline Free Choice data, the TC-A Free Choice data, and the TC-AB Free Choice data.

**Figure S6:**
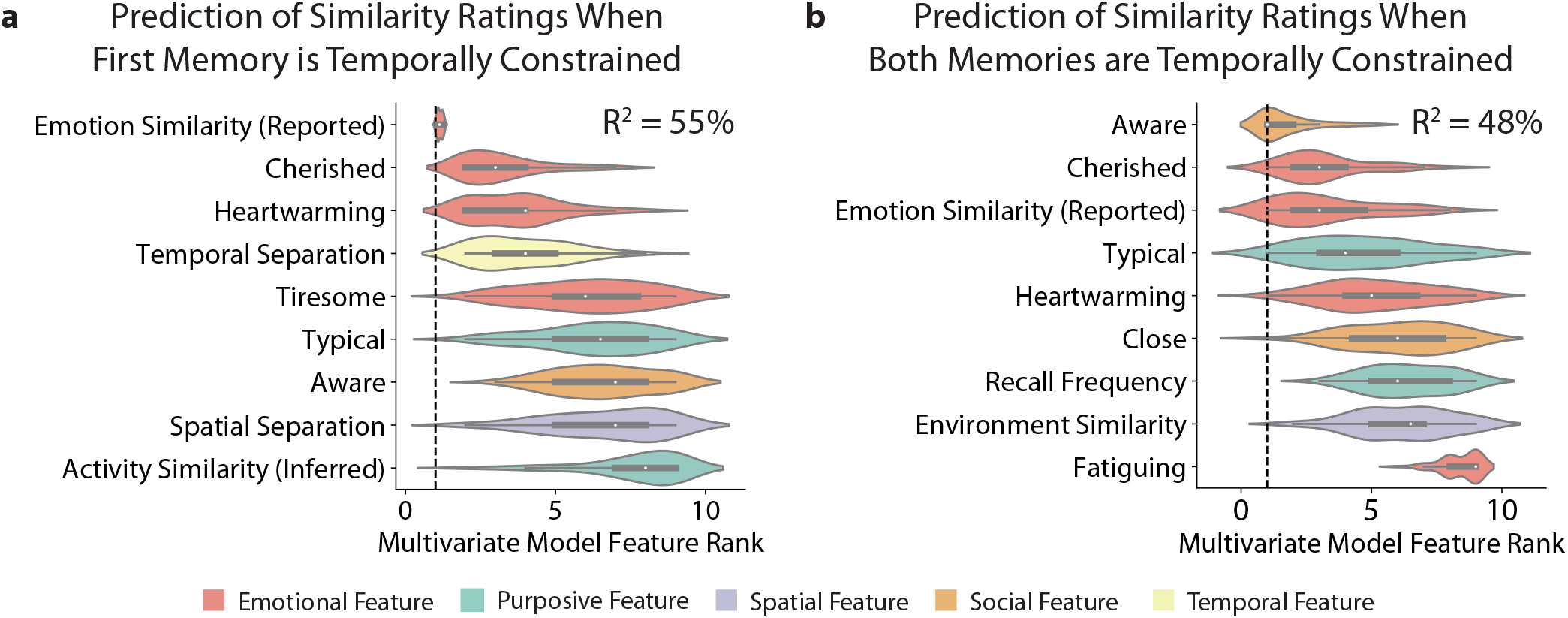
Multivariate regression analysis of the Temporally Constrained Free Choice condition. **(a):** Feature importance ranks of the features selected in the multivariate regression model when Memory A is constrained to be from a random two-hour time window the day before (TC-A). **(b):** The same as **(a)**, but analysis restricted to memory pairs for which Memory B came from the past week (TC-AB).

**Figure S7:**
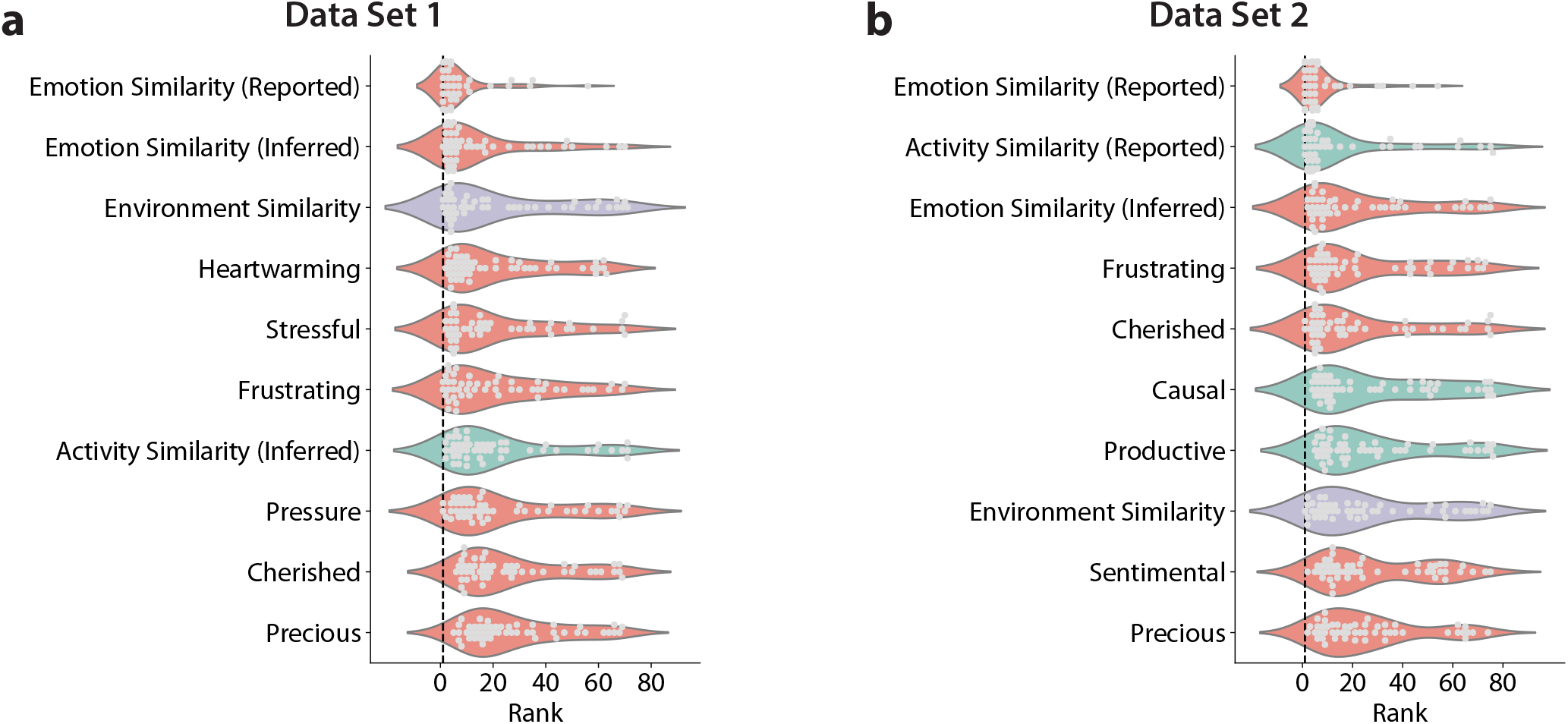
Distributions of univariate predictor relative rankings for each cross-validation fold. **(a):** Distributions of ranks for the top ten individual predictors of overall memory pair similarity in the original Free Choice data set, sorted from smallest median rank (best) to largest (worst). **(b):** The same as **(a)**, but for the second data set collected at a later time. The ordering of features according to median rank are largely consistent between Data Sets 1 and 2. In particular, Emotion Similarity (Reported) remains the strongest predictor of overall memory pair similarity.

**Figure S8:**
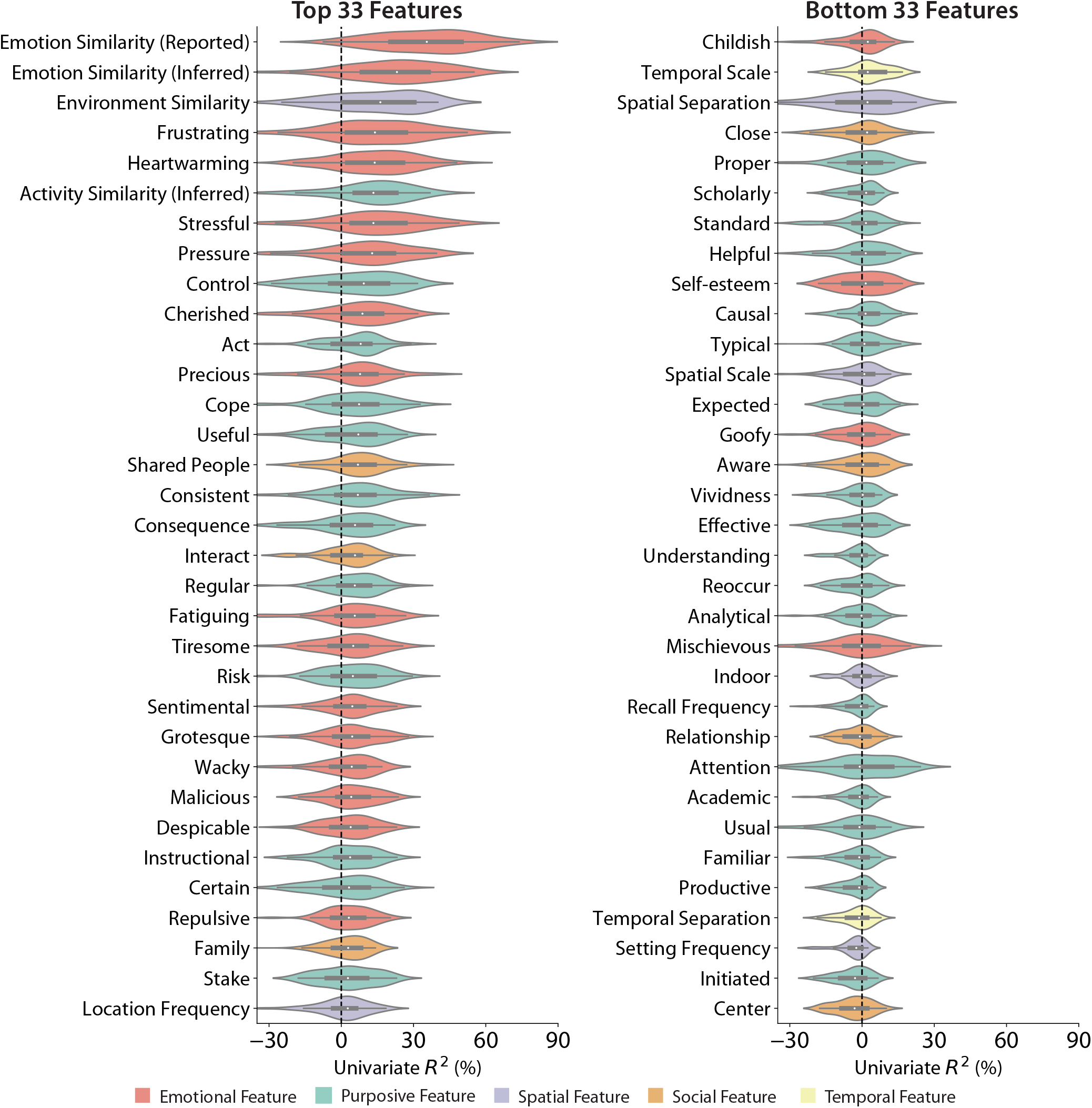
Free Choice condition distributions of *R*^2^ values for univariate regression of each feature on Overall Similarity. Features are sorted from greatest median *R*^2^ to least. This is the same as Figure 3f, but showing the full set of 66 features.

**Figure S9:**
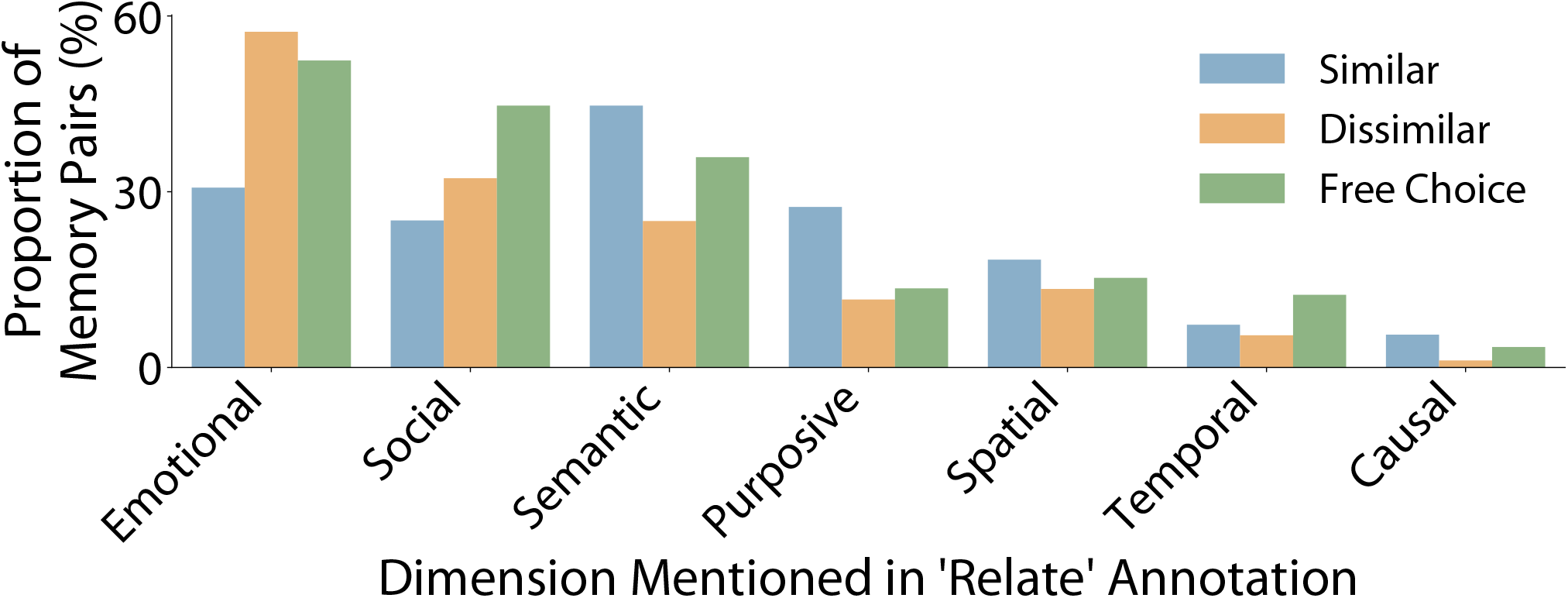
The proportion of participants who explicitly mentioned emotional, social, semantic, purposive, spatial, temporal, or causal/narrative-like reasons for why memory pairs were perceived as similar and/or dissimilar in the “Relate” annotation. Immediately after providing Memory A and Memory B, and before providing any memory annotations or ratings, participants were asked what made the memories similar (in the Similar Condition) or different (in the Dissimilar condition) or either (in the Free Choice condition); see Table S1, “Relate” annotation. A preliminary analysis of participants’ freely generated responses was performed by manually scoring their free responses. Free responses were scored by a single rater for their mentioning emotional, social, purposive, spatial, temporal or causal features. An example of an emotional reason would be “What makes these two memories similar is that both were happy.” A social reason would be “Memories A and B are dissimilar because in one I was with my husband while in another I was with a coworker.” A semantic reason would be “Memories A and B are similar because they were both about romance.” A purposive reason would be “Memories A and B are different because one was about watching a movie and the other was about completing a task at work.” A spatial reason would be “Memories A and B are similar because they both took place in parks.” A temporal reason would be “What makes Memories A and B dissimilar is that one was when I was around eight years old and the other was when I was in my late thirties.” A causal reason would be “What makes memories A and B similar is that the event in Memory B naturally led to the event in Memory A.”

**Figure S10:**
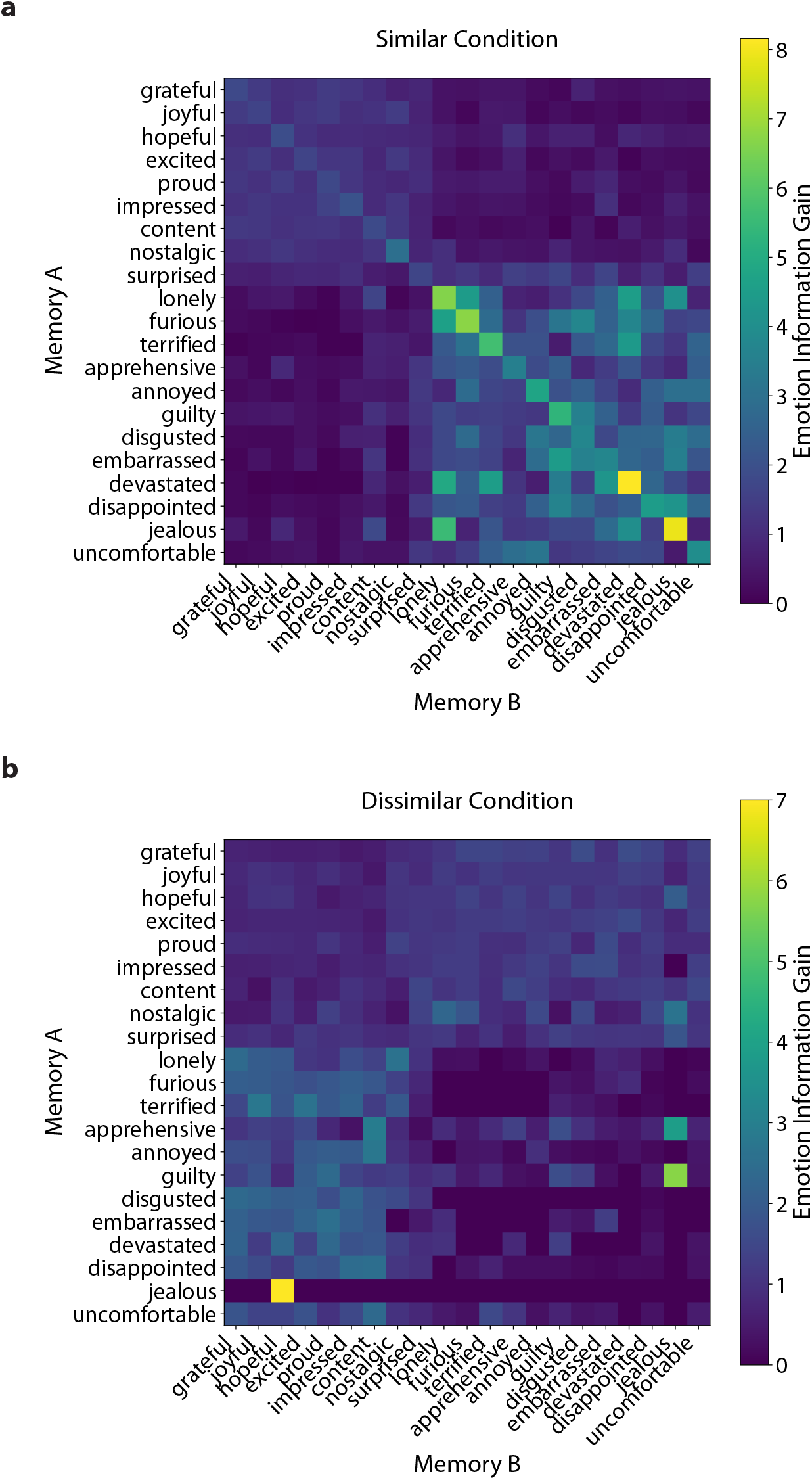
Memory pairs from the Similar condition tend to have shared emotional content, while memory pairs from the Dissimilar condition tend to have different emotional content. **(a):** Information gain of Memory B emotion when Memory A emotion is known, for memory pairs generated in the Similar condition. Specifically, if *P* (*A* = *i*) is the probability that Memory A is assigned emotion label *i* and *P* (*B*=j) is the probability that Memory B elicited from the same participant has label *j*, then the information gain, *I*(*i, j*) = *P* (*A* = *i B* = *j*)*/P* (*A* = *i*) = *P* (*A* = *i, B* = *j*)*/P* (*A* = *i*)*P* (*B* = *j*). Thus, information gain quantifies the change in probability of the combination of labels, over and above the joint probability they would have if they were independent. Emotions along the axes are clustered by affective valence. The noticeable diagonals indicate that memory pairs from the Similar condition tend to have the same specific emotional quality. The noticeable blocks on the top left and bottom right indicate that even when the specific emotional quality is not shared between Memory A and Memory B, there is still a tendency for them to share the same valence. **(b):** The same as (a), but for memory pairs generated in the Dissimilar condition. The noticeable blocks on the bottom left and top right indicate that memories from the Dissimilar condition tend to have different valence.

**Table. S1:**
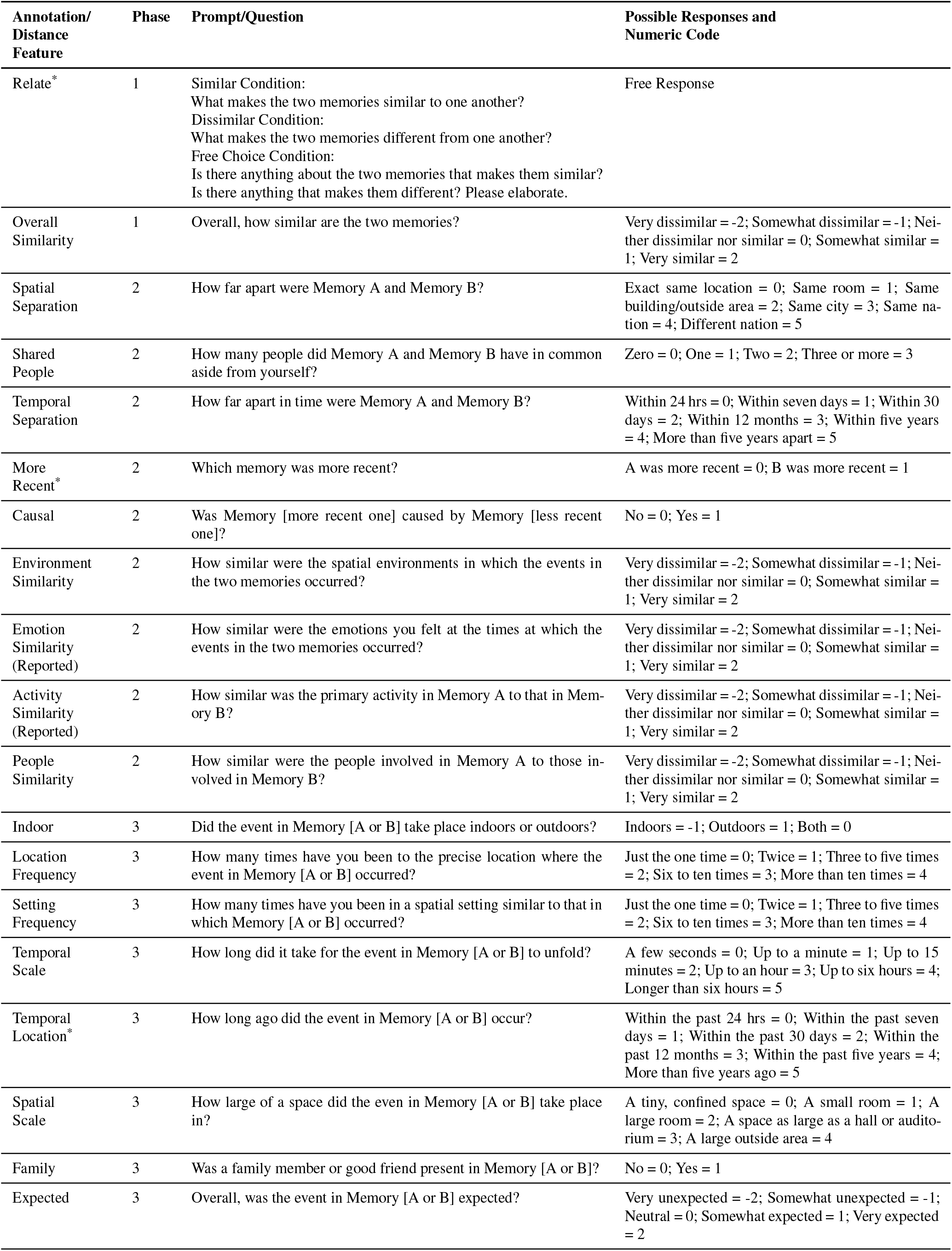
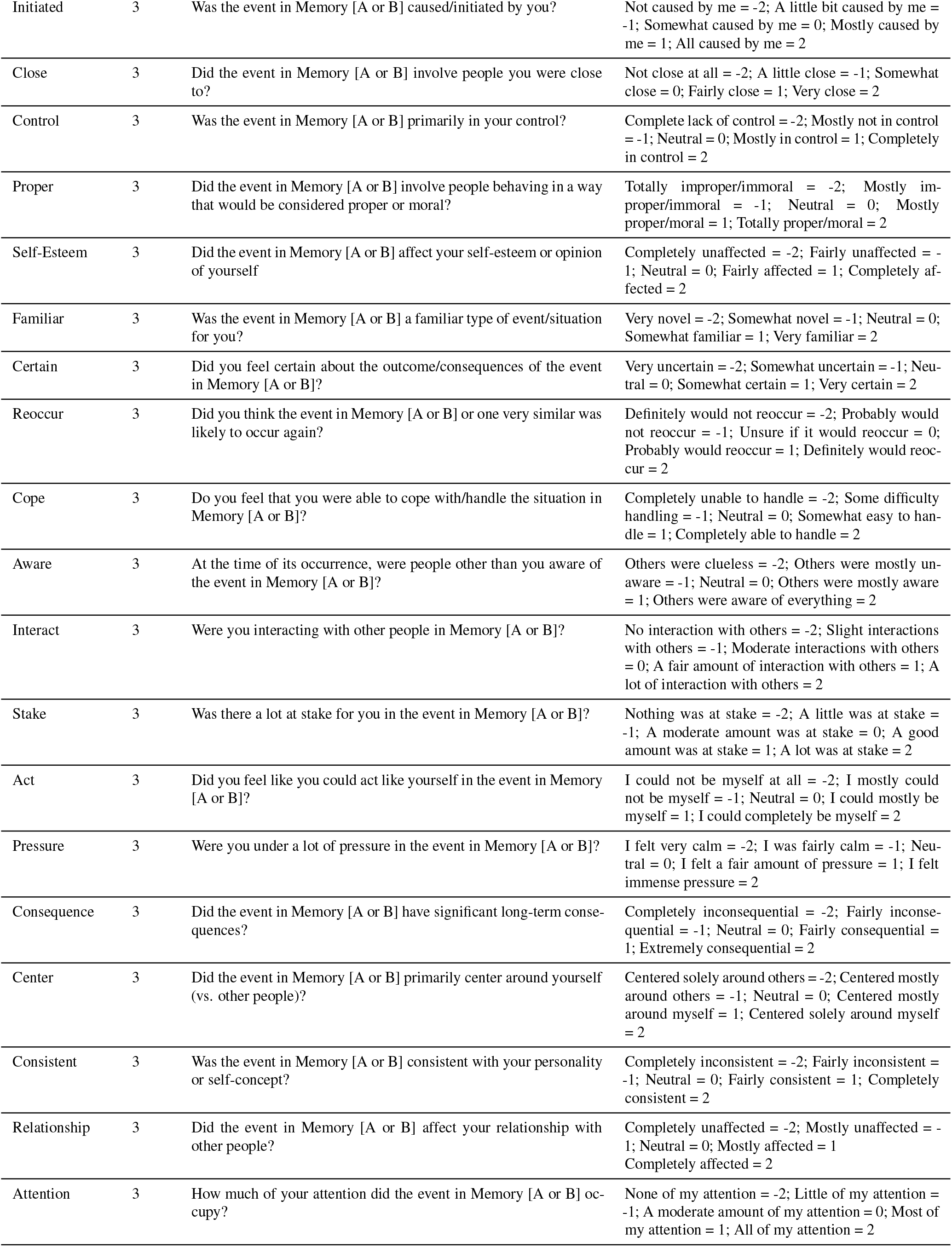
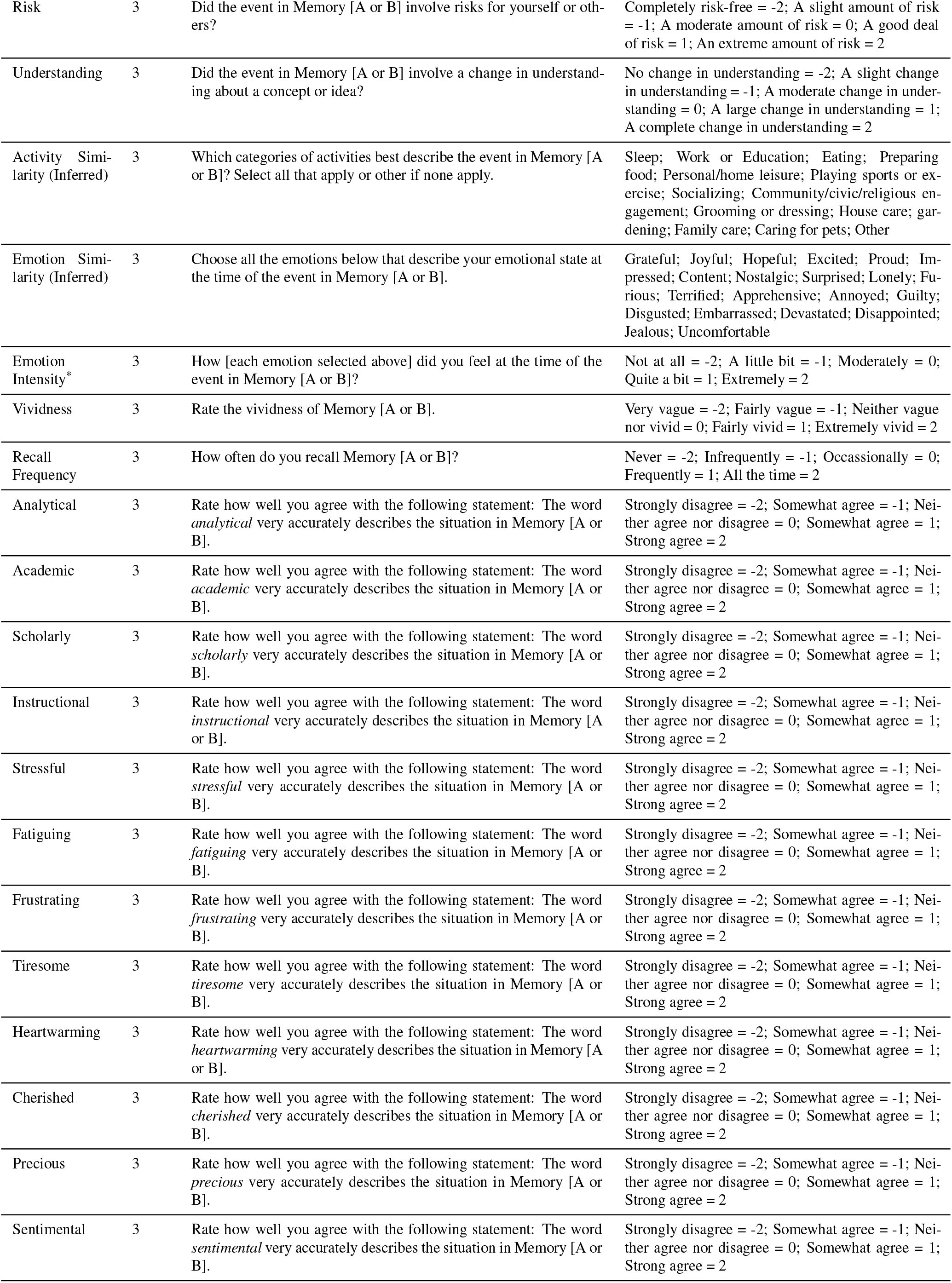
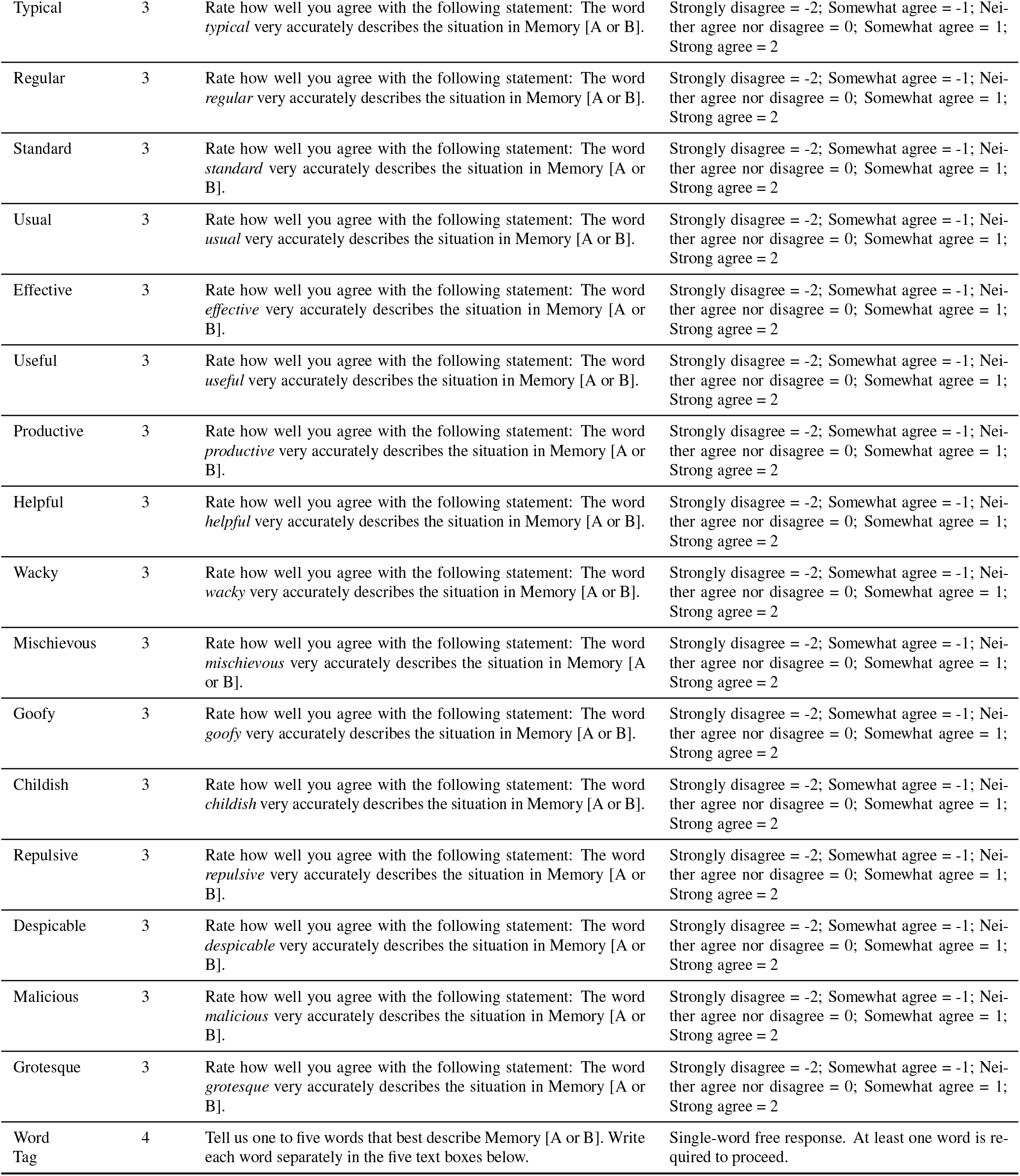
Features derived from the memory annotation stage of the experiment. ^*^ indicates annotations that were not used to derive a corresponding distance feature.

**Table. S2:** Summary of participants recruited and excluded from analysis. First participants were excluded based on insufficient catch trial performance (Criterion 1). Additional participants from the Similar and Dissimilar conditions were then excluded if their Overall Similarity rating was incongruent with the assigned condition (Criterion 2). From the remaining pool, participants were excluded if textual responses during the memory elicitation stage were manually identified as not being autobiographical memories (Criterion 3).

| Experiment | Condition | No. Recruited | No. Excluded -<br>Criterion 1 | No. Excluded -<br>Criterion 2 | No. Excluded -<br>Criterion 3 | No. Included |
| --- | --- | --- | --- | --- | --- | --- |
| Baseline | Similar | 224 | 21 | 14 | 10 | 179 |
|  | Dissimilar | 201 | 15 | 17 | 5 | 164 |
|  | Free Choice<br>(Data Set 1) | 198 | 16 | N/A | 12 | 170 |
|  | Free Choice<br>(Data Set 2) | 137 | 32 | N/A | 6 | 99 |
| Temporally<br>Constrained | Free Choice | 132 | 8 | N/A | 4 | 120 |

